# Dietary polyphenols drive alterations in behavior, transcriptional regulation, and commensal microbiota in models of opioid use

**DOI:** 10.1101/2022.06.14.496180

**Authors:** Aya Osman, Rebecca S. Hofford, Katherine R. Meckel, Yesha A. Dave, Sharon M. Zeldin, Ava L. Shipman, Kelsey E. Lucerne, Kyle J Trageser, Tatsunori Oguchi, Giulio M. Pasinetti, Drew D. Kiraly

## Abstract

Opioid Use Disorder (OUD) is a neuropsychiatric condition associated with tremendous medical and social consequences. Despite this burden, current pharmacotherapies for OUD are ineffective or intolerable for many patients. As such, interventions aimed at promoting overall health and resilience against OUD are of immense clinical and societal interest. Recently, treatment with a Bioactive Dietary Polyphenol Preparation (BDPP) was shown to promote behavioral resilience and adaptive neuroplasticity in multiple models of neuropsychiatric disease. Here, we assessed effects of BDPP treatment on behavioral and molecular responses to repeated morphine treatment. We find that BDPP pre-treatment alters responses across the dose range for both locomotor sensitization and conditioned place preference. Most notably, polyphenol treatment consistently reduced formation of preference at low dose (5mg/kg) morphine but enhanced it at high dose (15mg/kg). In parallel, we performed transcriptomic profiling of the nucleus accumbens, which again showed a dose x polyphenol interaction. At high dose morphine, BDPP pre-treatment potentiated gene expression changes induced by morphine particularly for genes related to synaptic function. We also profiled microbiome composition and function, as polyphenols are metabolized by the microbiome and can act as prebiotics. The profile revealed polyphenol treatment markedly altered microbiome composition and function, particularly in the low dose morphine group. Finally, we investigated involvement of the SIRT1 histone deacetylase, and the role of specific polyphenol metabolites in these behavioral phenotypes. Taken together, these results demonstrate that polyphenols have robust dose-dependent effects on behavioral and physiological responses to morphine and lay the foundation for future translational work.

## Introduction

Opioid use disorder (OUD) is a severe neuropsychiatric condition characterized by cycles of out of control drug intake, persistent use despite negative consequences, and cycles of abstinence and relapse [1,2]. Among substance use disorders, OUDs are particularly problematic given the high propensity for overdose [3,4]. Despite the consequences, rates of pathological OUD have continued to increase over the past decade, with early data suggesting the Covid-19 pandemic has exacerbated the severity of this epidemic [5]. There are currently multiple FDA-approved medications for the treatment of OUD, including opioid agonist replacement therapies and opioid receptor antagonists. While these therapies are effective for some, for many they are ineffective or intolerable [6]. Given the difficulties in treating patients once they have developed an OUD, developing interventions that can prevent the progression to pathological opioid use is of high priority in the field.

Opioid use disorder is widely believed to be the result of a constellation of neuroadaptations that occur through repeated or prolonged drug use [7]. These changes in epigenetic and transcriptional regulation result in altered functional and structural synaptic plasticity [8,9]. More recently, alterations in systemic inflammation and gut-brain signaling have been implicated in the maladaptive behavioral and synaptic plasticity resulting from prolonged opioid exposure [10,11].

Dietary polyphenols are a class of naturally occurring compounds from botanical sources which are neuroprotective in numerous models of neuropsychiatric disease [12–15]. In a stress-induced model of depression, treatment with polyphenols resulted in increased resilience to the formation of depression-like behaviors [16]. Similar effects have been found in models of neurodegenerative diseases [12,17]. In models of substance use disorders, treatment with dietary polyphenols has been linked to reduced formation of alcohol preference [18–20] and has been shown to decrease maladaptive neuroplasticity following treatment with psychostimulants [21–23]. While currently there is no published literature on polyphenols and OUD, there is significant literature showing polyphenols can reduce pain in models of pathological pain [24–28].

Polyphenol compounds exert their neuroprotective effects via myriad mechanisms. They are well known to reduce inflammation and alter redox balance [16,29,30]. Polyphenol compounds, most notably resveratrol, activate the sirtuin family of histone deacetylases, leading to altered epigenetic regulation of gene expression and altered neurobiological and behavioral plasticity [31]. Additionally, polyphenols also exert effects on transcriptional control of gene expression and neuronal function via sirtuin-independent mechanisms [32–34]. Additionally, polyphenols extensively interact with the gut microbiome in ways that are critical for gut-brain signaling [35].

Here, we use the previously described brain-penetrant bioactive dietary polyphenol preparation (BDPP) [16,32] to assess how polyphenols may influence the formation of addiction-like behaviors and transcriptional regulation in models of OUD. We find that pre-treatment with BDPP can reduce the formation of morphine locomotor sensitization and conditioned place preference. Additionally, we find that treatment with polyphenols interacts with morphine to alter the composition of the gut microbiome and the transcriptome in the nucleus accumbens. Taken together, these findings suggest translational potential for polyphenols in reducing the formation of opioid addiction-like behaviors.

## Materials & Methods

### Animals

Male C57BL/6J mice (7-9 weeks old, Jackson Laboratories) were group-housed (4-5 mice/cage) in a humidity and temperature-controlled colony room on a 12/12h light-dark cycle (lights on at 7:00am). Drink solutions and food were available ad libitum throughout the entirety of all experiments. All animal procedures were approved by the Mount Sinai IACUC and all procedures conformed to the “Guide for the Care and Use of Laboratory Animals” (National Research Council 2010).

### Preparation and delivery of dietary polyphenols

Cages were randomly assigned to control or BDPP treatment. BDPP treatment consisted of 100 ml of Concord (Welch’s grape juice), 0.4g Grape Seed Polyphenolic Extract (GSPE) (Healthy Origins #57914), and 0.4g resveratrol (Bulk supplements #SKU RES100GC) in 300 ml water as described previously [16,32]. Control animals were provided with water containing matched sucrose content. Mice were treated for 2 weeks prior to the start of experimental procedures and all mice remained on their drink solutions until the conclusion of the studies.

### Morphine

Morphine sulfate was provided by the NIDA drug supply program from National Institute on Drug Abuse and was diluted in saline and injected subcutaneously.

### Locomotor Sensitization

Acute and sensitized locomotor responses to morphine were carried out using San Diego Instruments Photobeam Activity System, largely as described previously. Details available in **Supplemental Methods**.

### Morphine conditioned place preference

Mice from Control and BDPP treatment groups were injected with 2.5, 5, or 15mg/kg morphine and underwent morphine conditioned place preference (CPP) largely as described previously [36]. Detailed methods in **Supplemental Methods**.

### Withdrawal study CPP

Mice were injected with 5mg/kg morphine in their home cages for 5 consecutive days. They were then given control or BDPP treatment for two-weeks during drug abstinence. Mice then underwent CPP as described above to measure the effect of morphine pre-exposure on formation of preference.

### RNA-Sequencing of NAc

Control or BDPP treated mice underwent CPP paradigm injected with saline, 5mg/kg, or 15mg/kg morphine and were rapidly decapitated 1hr post-test session of CPP. The Nucleus Accumbens (NAc) was flash frozen on dry ice, and RNA isolation, Assessment of RNA integrity, Poly-A library preparation, and 2×150bp paired-end sequencing were performed according to standard procedures – details in **Supplemental Methods**.

### RNA-seq Data Analysis

For differential expression analysis, aligned reads were normalized to Log2 counts per million, and differential expression was analyzed using Deseq2 with statistical significance set at a threshold of FDR-corrected *p* < 0.2 except as described. Identification of significantly enriched gene ontologies and transcription factors was performed using g:Profiler and Enrichr software packages [37–39]. Further details in **Supplemental Methods**.

### 16s-Sequencing of Cecal Content

Control or BDPP treated mice underwent CPP paradigm injected with 5mg/kg or 15mg/kg morphine and were euthanized 24h post-test session. The caecum was rapidly dissected, contents flash frozen on dry ice and stored at -80C. DNA isolation and sequencing were carried out using standard procedures. Data analysis was performed using the QIIME and PICRUSt2 analysis packages with standard settings [40,41] – full details in **Supplemental Methods**.

### SIRT1 Inhibitor Study

To assess the role of SIRT1 in mediating BDPP effects, control or BDPP treated mice were anesthetized with a combination of ketamine (100mg/kg) and xylazine (10mg/kg) and surgically implanted with bilateral cannula targeting the NAc under stereotactic guidance. The coordinates from bregma were: anteroposterior 1.5mm, mediolateral 1.0mm, dorsoventral 4.5 mm. Animals were allowed to recover from surgeries for one week prior to undergoing CPP as described above using 5mg/kg morphine. Animals were maintained on drink solutions throughout. On each of the first four days of CPP 0.05mM of EX-527 (Sigma-Aldrich) a SIRT1 specific inhibitor or vehicle (DMSO) was infused into the NAc over 5 mins immediately after the morphine session.

### DHCA/Mal-Gluc Treatment Study

Mice were randomly divided into two groups: control group provided with standard drinking water, and a group treated with a mixture of dihydrocaffeic acid (**DHCA** - 5mg/kg-BW/day) and malvidin-3′-O-glucoside (**Mal-gluc** 0.5 µg/kg/day), delivered through their drinking water as previously described [16], both treatments starting 2 weeks prior to CPP using 5mg/kg and 15mg/kg morphine.

### Statistical analysis and Figures

All behavioral analyses were performed using GraphPad Prism with two-way ANOVAs with repeated measures as appropriate for 2 × 2 experiments, and as two-tailed T-tests for pairwise comparisons. 16S and RNA sequencing data were analyzed as detailed in **Supplemental Methods**. Graphs of all figures were created in Graphpad Prism and R. Experimental timelines were generated in BioRender with full permission to publish.

## Results

### Controls for Polyphenol Treatment

For most studies C57BL/6J mice were treated with BDPP polyphenol cocktail or control water in their home cage two weeks prior to the start of experimentation and remained on these drink solutions throughout duration of the study (**Fig. 1A**). A subset of mice were monitored for water intake and bodyweight change as indicators of overall health. We found no significant difference in drink intake (**Fig. 1B** - *t*_(35)_=1.45, *p*=0.16) or bodyweight change between the groups (**Fig. 1C** - *t*_(30)_=0.15, *p*=0.88).

**Figure 1.**
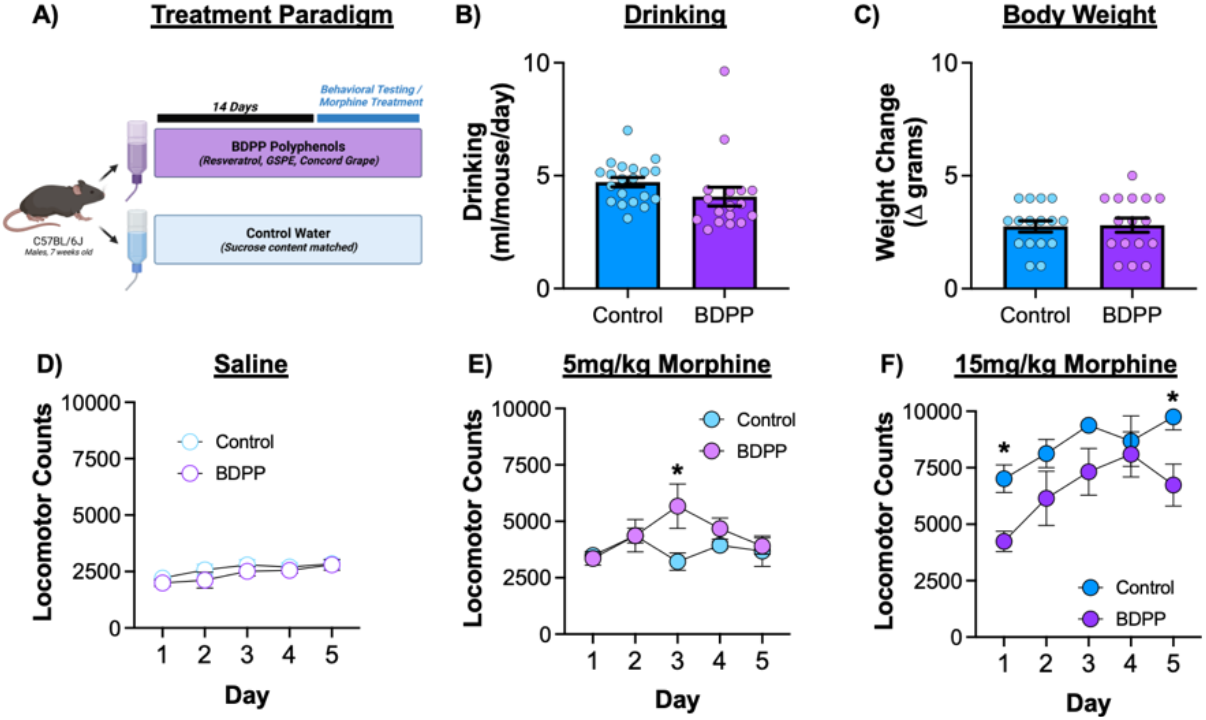
Baseline effects of BDPP polyphenols and locomotor response to morphine. **(A)** Graphical timeline of the associated studies. **(B)** To ensure that treated water did not affect drinking we measured consumption in a subset of animals during the first two weeks. **(C)** Body weight change over the course of the experiment was also measured. **(D)** Repeated injections of saline did not affect locomotor activity in either treatment group. **(E)** Repeated 5mg/kg injections of morphine resulted in a significant time x treatment interaction with polyphenol animals having increased locomotion on day 3. **(F)** Polyphenol treatment resulted in decreased development of locomotor sensitization to 15mg/kg morphine. Data presented as means ± SEM. * *p* < 0.05; ** *p* < 0.01.

### Locomotor Sensitization

To determine the effects of BDPP supplementation on behavioral response to opioids, we measured activity in a locomotor sensitization paradigm. As a control we tested all animals with five daily repeated injections of saline. There was no effect of BDPP treatment (**Fig. 1D** – two-way RM-ANOVA: F_(1,14)_=1.30; *p*=0.27), a modest effect of time (F_(4,56)_=4.83; *p*=0.002) but no significant interaction (F_(4,56)_=0.36; *p*=0.83). At the lower 5mg/kg dose of morphine there was no effect of treatment (F_(1,11)_=1.39; *p*=0.26) or time (F_(4,44)_=2.20; *p*=0.085). However, there was a significant treatment by time interaction (F_(4,44)_=3.15; *p*=0.02), with post-hoc testing demonstrating significant increase in locomotor response in the BDPP group on Day 3 (**Fig. 1E**). Sensitization effects at 15mg/kg dose of morphine resulted in effects of both treatment (**Fig. 1F** - F_(1,10)_=12.49; *p*=0.005) and time (F_(4,39)_=4.3; *p*=0.006) with the BDPP group showing reduced locomotor activation in response to acute and repeated morphine. There was no significant time x treatment interaction (F_(4,39)_=0.68; *p*=0.6). Additional details on acute and sensitized locomotor response are available in **Supplemental Results** and **Fig. S1**.

### Conditioned Place Preference

While locomotor sensitization is an important marker of plasticity in response to drugs of abuse, it has little specificity for assessing rewarding drug effects. To test how BDPP pre-treatment modulated the rewarding effects of morphine we performed CPP testing for morphine (**Fig. 2A**). When assessed by two-way ANOVA we find that there was a trend but no effect of BDPP treatment (F_(1,114)_=3.31; *p*=0.07), and no effect of morphine dose (F_(2,114)_=1.35; *p*=0.26). However, there was a BDPP x dose interaction (F_(2,114)_=8.17; *p*=0.0005) – which is likely responsible for abrogating the main effect of dose. On post-hoc testing, BDPP treatment led to a marked decrease in preference at the intermediate 5mg/kg dose (**Fig. 2B middle** – *p*=0.002), but a significant increase in CPP at 15mg/kg (**Fig. 2B right** – *p*=0.03).

**Figure 2.**
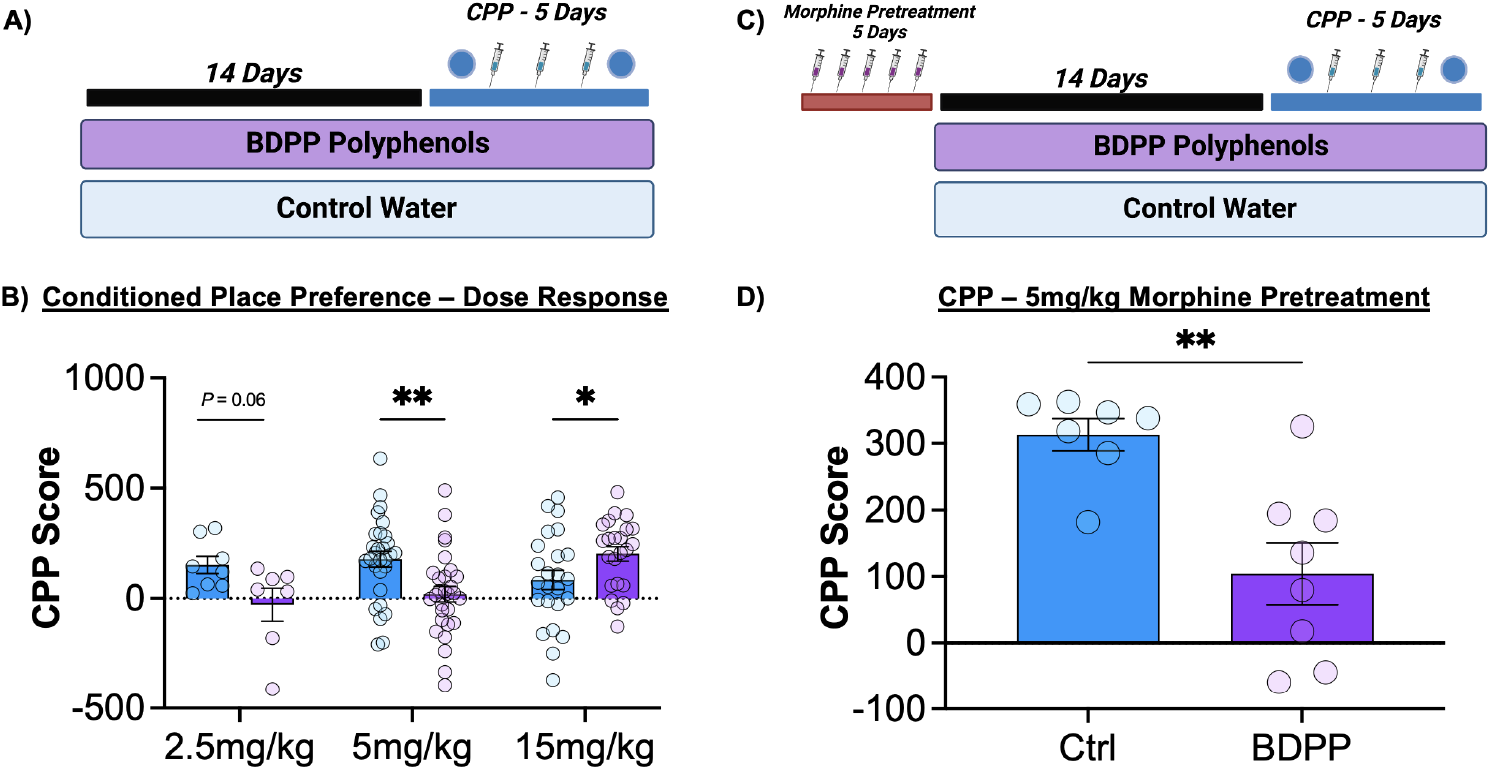
Effects of BDPP Polyphenols on formation of morphine conditioned place preference. **(A)** After two weeks of BDPP or control pretreatment, mice were tested on a conditioned place preference assay using a five-day protocol. **(B)** CPP results across a dose range showed a robust dose x treatment interaction with BDPP-treated mice showing decreased preferences at lower doses of morphine, but increased preference for the higher 15mg/kg dose. **(C)** In a subsequent experiment, mice were pretreated with morphine prior to polyphenol treatment and CPP testing. **(D)** While morphine pretreatment enhanced subsequent formation of CPP, BDPP polyphenol pretreatment still resulted in reduced formation of preference at 5mg/kg morphine. Data presented as means ± SEM. * *p* < 0.05; ** *p* < 0.01.

As prior treatment with opioids can lead to potentiation of future behavioral response in subsequent place preference [42], we then tested the effect of BDPP on CPP for intermediate dose morphine using an opioid re-exposure model. Mice received injections of 5mg/kg morphine for five days prior to the start of polyphenol treatment, they then underwent the normal BDPP treatment and CPP procedure (**Fig. 2C**). As predicted, the pre-treatment with morphine potentiated morphine CPP compared to the standard regimen (**Fig. S2**). However, even when mice had been pre-treated with morphine prior to polyphenols, the two weeks of BDPP again resulted in a reduction of CPP (**Fig. 2D** – t_(13)_=3.81; *p*=0.002).

### RNA-Sequencing of the Nucleus Accumbens

Substance use disorders and behavioral adaptations to drugs of abuse are dependent on alterations in regulation of gene expression [8]. Additionally, dietary polyphenols, alter gene expression in the brain [16,43]. To better understand how BDPP treatment alters neurobiological response to morphine, we performed RNA-sequencing of the nucleus NAc – a key structure in driving behavioral response to opioids [44]. For these experiments, animals underwent CPP as normal, and a control group received saline injections during each session (**Fig. 3 – top**).

**Figure 3.**
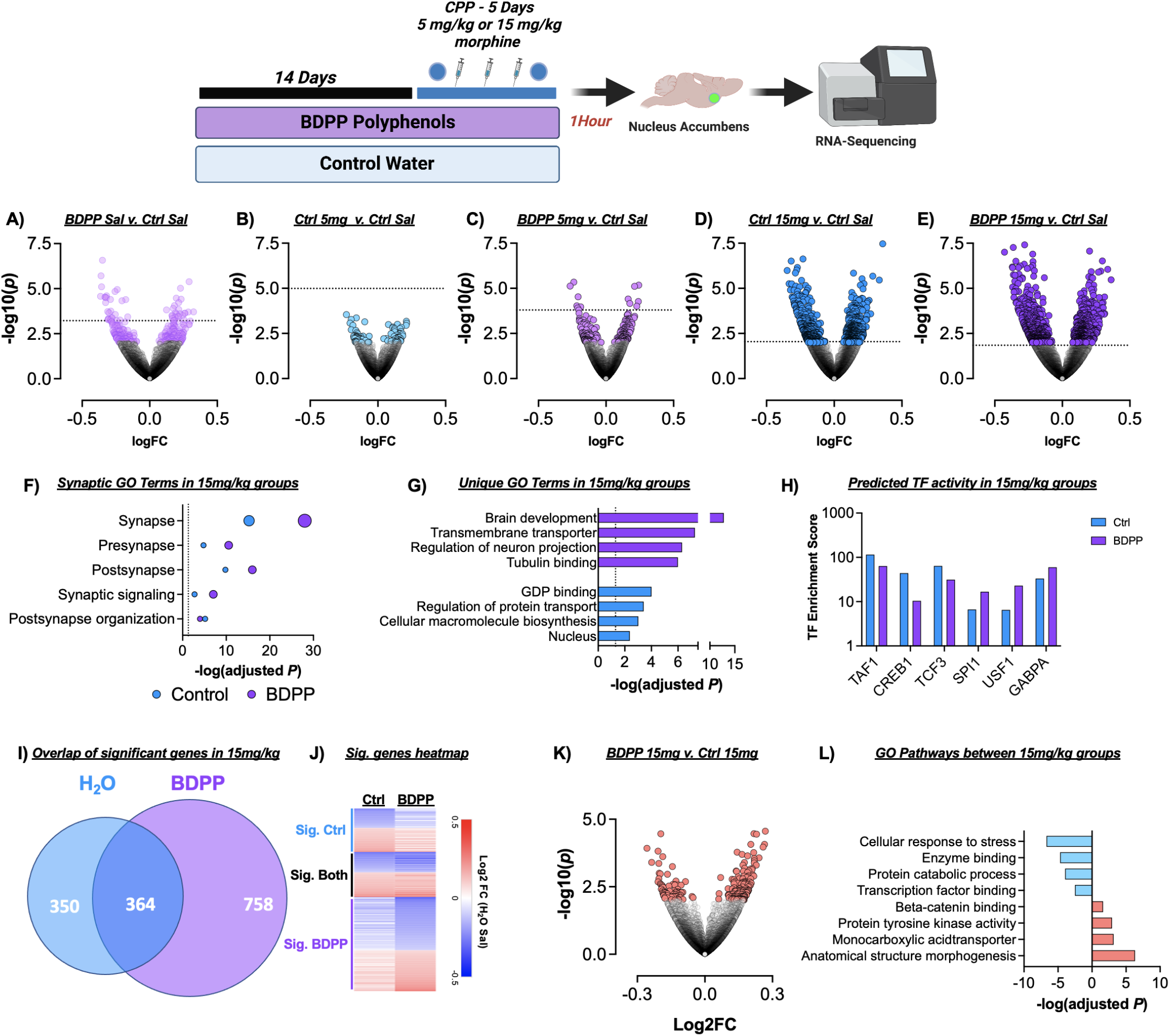
Effects of BDPP polyphenol treatment on transcriptional response to morphine in the nucleus accumbens. **(Top)** Timeline for sequencing experiments. Mice were pretreated for two weeks prior to CPP training and were sacrificed one hour after the post test session. **(A-E)** Volcano plots of all treatment groups relative to Control Saline group. **(F)** Gene ontology analysis of synapse related genes in H2O and BDPP 15mg/kg groups. **(G)** GO terms uniquely regulated in BDPP 15mg/kg and Ctrl 15mg/kg groups compared to Ctrl Saline. **(H)** Transcription factor enrichment analysis of top enriched treatment groups in Ctrl and BDPP 15mg/kg groups. **(I)** Venn diagram off all genes statistically significant in Ctrl and BDPP 15mg/kg groups relative to Ctrl Saline. **(J)** Heatmap of fold change expression of all genes from the previous panel. **(K)** Volcano plot of all significantly different genes between the two morphine 15mg/kg groups. Fold change is relative to Ctrl 15mg/kg. **(L)** Gene ontology analysis of top terms regulated up or down between the two 15mg/kg morphine groups.

To obtain a broad understanding of differential gene expression, we created volcano plots of all groups compared to Control-Saline (**Fig 3 A-E**, colored circles *p*<0.01). As expected, the most robust effects on gene expression were seen with the higher 15mg/kg dose of morphine (**Fig. 3 D,E**). Interestingly, treatment with BDPP resulted in more significantly regulated genes at both doses, and treatment with BDPP alone resulted in more significantly regulated genes than mice treated with water or BDPP in combination with 5mg/kg morphine (**Fig. 3 A-C**). Full differential gene lists for each pairwise comparison available as **Supplemental Tables 1-5**.

We then examined changes in genes that met statistical significance (FDR *p*<0.2) (**Fig. 3 A-E** dotted line indicates the nominal *p* value at which this threshold is met). As there were more significantly regulated genes in the 15mg/kg morphine groups, (**Fig. 3 D,E**) detailed pathway analysis was conducted for the 15mg/kg groups. Both control and BDPP 15mg/kg morphine treated mice had significant enrichment of genes related to the synapse, both pre and post synaptic function, synaptic signaling, and postsynaptic organization when compared to control Saline (**Fig. 3F;** Full GO **Tables S6-7**). In all these pathways enrichment was more robust in the BDPP 15mg/kg morphine animals.

Additionally, there were numerous uniquely enriched pathways in the two groups. BDPP 15mg/kg mice showed significant enhancement of brain development, transmembrane transporters, and tubulin binding (**Fig. 3G**), while control 15mg/kg mice had enrichment of GDP-binding, macromolecule biosynthesis, and nuclear genes. Next, the Enrichr software package was used to analyze differences in transcription factor involvement between control 15mg/kg and BDPP 15mg/kg morphine groups. We find that treatment with control or BDPP resulted in differential recruitment of transcription factors (**Fig. 3H** – note log10 y-axis; full Enrichr results **Table S8**).

Of the 1472 genes that were regulated compared to control saline between the two 15mg/kg morphine groups, 364 were significantly regulated in both groups, while 350 uniquely changed in control and 758 uniquely changed in BDPP (**Fig. 3I**). When all of these genes were visualized as a heatmap examining fold change relative to control Saline animals, nearly all genes change in the same direction (**Fig. 3J**) –but not necessarily statistically significant in both.

Finally, both 15mg/kg morphine groups were compared to examine how the addition of BDPP pre-treatment affected gene expression when compared directly to other morphine paired animals. **Figure 3K** shows a volcano plot depicting the 205 genes that were statistically significant at a *p*<0.01 threshold (67 downregulated, 139 upregulated) between control 15mg/kg and BDPP 15mg/kg. Gene ontology analysis was performed separately on the up and downregulated gene lists. BDPP 15mg/kg mice showed decreased expression in pathways related to cellular response to stress and transcription factor binding among others, and increases in pathways related to anatomical structure, morphogenesis, and protein tyrosine kinase activity (**Fig. 3L** – Full table: **Table S9**).

### 16s Sequencing of Cecal Content

Given the interactions between dietary polyphenols and the microbiome [45,46], effects of BDPP treatment and interactions with morphine were assessed (**Fig.4A**). Analysis of alpha diversity, which calculates richness and evenness of microbial species, using the Chao1 metric [47] showed a significant effect of BDPP (**Fig. 4B** - F_(1,28)_=5.06; *p*=0.03), no effect of morphine dose (F_(1,28)_=1.05; *p*=0.31) and no significant BDPP x morphine dose interaction (F_(1,28)_=0.82; *p*=0.37). Post-hoc testing showed BDPP treatment led to significantly decreased alpha diversity only in the 5mg/kg morphine group (**Fig. 4B – left** – *p*=0.03). Next β-diversity, a measure of between-subject diversity, was assessed which revealed a clear separation between control and BDPP treated animals in the 5mg/kg morphine group. However, no separation between water control and BDPP treated animals was observed in the 15mg/kg morphine group (**Fig. 4C**).

**Figure 4.**
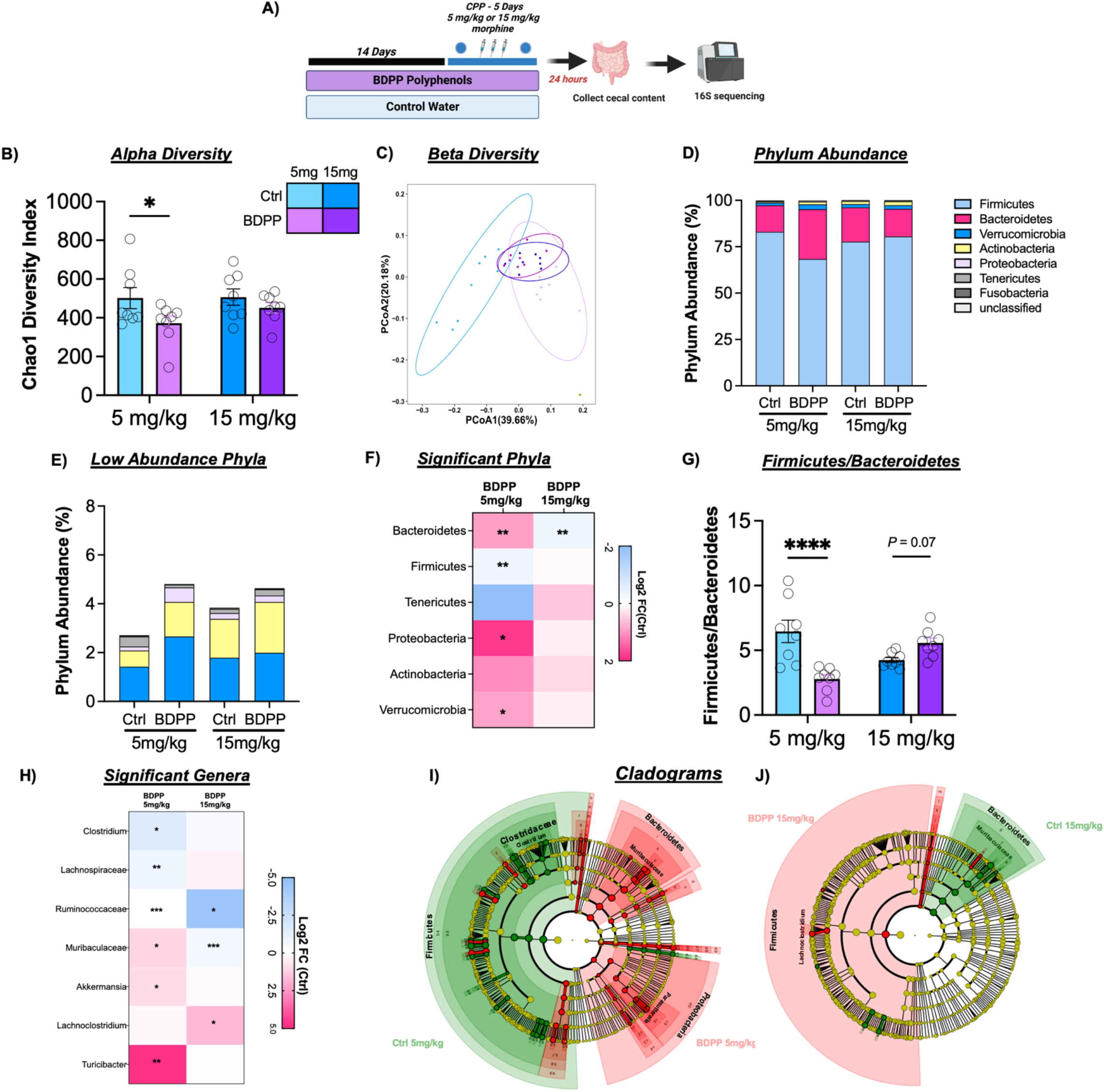
Effects of BDPP treatment on cecal microbial composition in morphine treated mice. **(A)** Timeline for 16s sequencing experiment. Mice were pre-treated for two weeks prior to CPP training and were sacrificed 24 hours after the post test session. **(B)** Alpha diversity of the gut microbiomes were calculated using Chao1 diversity metric, and shows reduced Alpha diversity in 5mg/kg BDPP treated mice **(C)** Beta diversity measured using the unweighted Unifrac distance metrics and shows BDPP treated mice possess a unique microbiome from control treated mice in 5mg/kg morphine group, but overlap in 15mg/kg morphine group **(D)** Stacked bar chart showing the relative phylum abundance in mice from all treatment groups, each phyla represented in a different color **(E)** Stacked bar chart showing phyla expression levels of the low abundance phyla (this chart corresponds to the top section of the graphs from panel D). **(F)** Heatmap showing changes in phylum diversities with BDPP treatment relative to controls at each dose. Asterisks represent *p* values from Wilcoxon text. **(G)** BDPP treated mice show a significant decrease in Firmicutes to Bacteroidetes Ratio compared to controls in the 5mg/kg morphine treatment group **(H)** Heatmap displaying the log2 Fold Change (FC) of selected altered bacterial genera in BDPP treated animals relative to respective control counterparts in 5mg/kg and 15mg/kg morphine group. Asterisks represent *p* values from Wilcoxon text. **(H)** Cladogram representing taxonomic biomarkers characterizing the differences between BDPP and control treated mice in the 5mg/kg morphine group **(I)** Cladogram representing taxonomic biomarkers characterizing the differences between BDPP and control treated mice in the 5mg/kg and 15mg/kg morphine groups (Full Key for figures **H** and **I** found in Tables **S13**). \**p* < 0.05 \*\**p* < 0.01 \*\*\**p* < 0.001.

Next, the effect of BDPP treatment on phylum abundance was examined. In the qualitative stacked bar plots in **Figures 4 D&E**, we see that BDPP results in shifts in phylum expression in the 5mg/kg group in the highly abundant Bacteroidetes and Firmicutes phyla (**Fig. 4D** – pink and blue bars), as well as in the less abundant phyla (**Fig. 4E**). Given that in mammals Bacteroidetes and Firmicutes make up ∼90% of total bacteria found in the gut, the relative abundance of these phyla has thus been tied to the health of the microbiome [48]. In the 5mg/kg morphine group, BDPP treatment significantly decreased the abundance of bacteria from the Firmicutes phylum (**Fig. 4F left** – Wilcoxon rank-sum test: *p*=0.002) and significantly increased the abundance of Bacteroidetes (*p*=0.002). In the 15mg/kg morphine treatment group, the opposite effect is observed where BDPP treatment resulted in a trend towards an increase in the abundance of Firmicutes (**Fig. 4F right** - Wilcoxon rank-sum test: *p*=0.046; q=0.23) and a significant decrease in Bacteroidetes *(p*=0.006) compared to control animals. Full list of phyla changes and statistics in **Table S10**.

Shifts in the ratio of the Bacteroidetes and Firmicutes phyla can be an important marker of microbiome health and stability. When the ratio of these two major phyla is compared, there was a modest main effect of BDPP treatment (**Fig. 4G** – F_(1,28)_=5.34; *p*=0.03), and no effect of morphine dose (F_(1,28)_=0.30; *p*=0.59). There was, however, a robust dose x treatment interaction (F_(1,28)_=23.91; *p*<0.0001). Post-hoc testing comparing the treatment groups within each dose showed a significant effect of BDPP at 5mg/kg (**Fig. 4G left** – *p*<0.0001).

We next examined phylogenetic changes down to the genus level. In the 5mg/kg group, bacteria belonging to the genus *Clostridium* were significantly reduced in BDPP treated animals (**Fig. 4H** – Wilcoxon, *p*=0.02; Full **Table S11**) however this reduction was not observed in 15mg/kg morphine treated animals (*p*=0.17). In the 5mg/kg morphine group BDPP treatment resulted in a significant increase in a genus of *Muribaculaceae* (*p*=0.01), but this same genus was decreased at 15mg/kg morphine (*p*=0.0008). Effects of BDPP treatment on the gut microbiome at the genus level are observed more clearly in the low dose morphine group, with significant increases in beneficial genera *Parasutteralla* (*p*=0.01) and *Akkermensia* (*p*=0.02). This is in line with the body of literature linking polyphenol treatment with increases in beneficial bacteria [49,50].

Next, *LEfSe* analysis was used to determine treatment-defining expression patterns [51]. Full differential bacterial group lists for each pairwise comparison available as **Fig.S3A** and **S3B**. Cladograms show that after 5mg/kg morphine, BDPP treated mice were characterized predominantly by a Proteobacteria/*Parasutterella* and Bacteroidetes/*Muribibaculaceae* enterotype (**Fig. 4I** – **red**), while controls were characterized by a contrasting Firmicutes/Clostridium enterotype (**Fig. 4I green**). Interestingly, in the15mg/kg morphine groups cladograms show an opposite effect, where 15mg/kg BDPP treated mice are characterized predominantly by a Firmicutes/*Lachnoclostridium* enterotype (**Fig. 4J-red**). Further detailed analysis of functional microbiome groups and correlations with behavior are available in the **Supplemental Results, Figs. S4-5, & Tables S12-15**.

### Mechanistic interrogation of polyphenol effects

We finally performed two targeted experiments to refine our mechanistic understanding of how BDPP treatment alters behavioral response to morphine. For the first, we based our experiment on the hypothesis that resveratrol and other polyphenols are activators of the SIRT1 lysine deacetylase - which has been shown to be important for behavioral response to opioids [52]. For these experiments, mice were treated with control water or BDPP as previous but had the specific SIRT1 inhibitor infused directly into their NAc after each of the first four days of CPP (**Fig. 5A**). For this experiment we still found a main effect of BDPP treatment (**Fig. 5B** - F_(1,13)_=6.38; *p*=0.025), similar in magnitude and direction to what was seen with BDPP treatment alone (**Fig. 2B**). However, there was no main effect of EX527 treatment (F_(1,13)_=0.44; *p*=0.52) or significant BDPP x EX527 interaction (F_(1,13)_=0.02; *p*=0.9). This indicates that effects of BDPP on CPP behavior at 5mg/kg are not mediated by SIRT1 activity in the NAc.

**Figure 5.**
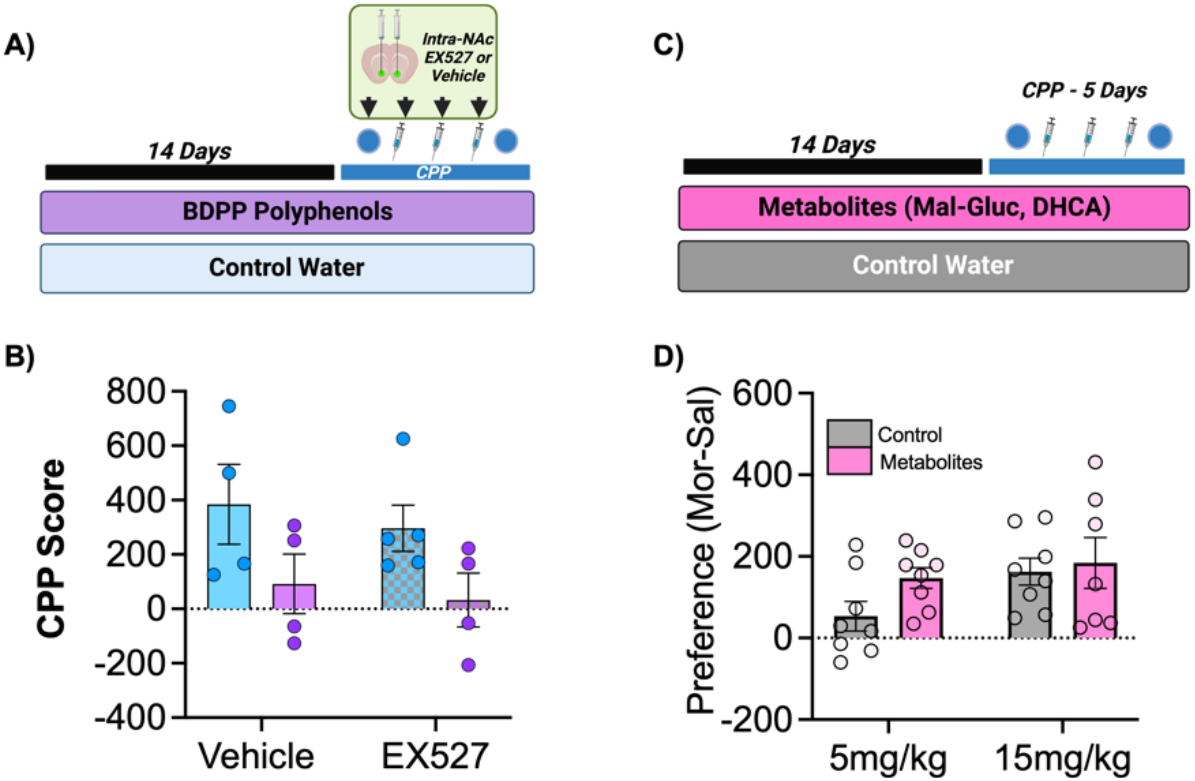
Mechanistic studies of polyphenol effects. **(A)** To test the contribution of SIRT1 activation to the behavioral effects of BDPP treatment a SIRT1 inhibitor was infused into the NAc during CPP. **(B)** CPP at 5mg/kg morphine again showed a main effect of BDPP treatment, but no effect of SIRT1 inhibition. **(C)** Effects of treatment with two key metabolites from the BDPP cocktail was assessed. **(D)** Treatment with BDPP metabolites did not result in significant changes in morphine preference at either dose.

We next examined if the effects of BDPP treatment could be explained by a subset of neuroactive metabolites derived from the mixture. Two metabolites from the BDPP cocktail, dihydrocaffeic acid (DHCA) and malvidin-3′-O-glucoside (Mal-gluc) were previously reported as key mediators of the behavioral and epigenetic effects of BDPP [16]. Thus mice were treated either with control water or Mal-gluc & DHCA treated water for two weeks prior to 5 and 15mg/kg morphine CPP (**Fig. 5C**). For these experiments there was no main effect of metabolite treatment (**Fig. 5D** - F_(1,27)_=2.1; *p*=0.16), a trend towards main effect of dose (F_(1,27)_=3.33; *p*=0.08), and no significant dose x treatment interactions (F_(1,27)_=0.81; *p*=0.38). These data demonstrate that the effects of the BDPP on addiction-like behaviours are not easily explained by a specific subset of BDPP derived metabolites and may require a more complex mixture of metabolites.

## Discussion

Here we demonstrate treatment with a bioavailable oral polyphenol mixture (BDPP) has marked effects on both the molecular and behavioral responses to morphine. Pretreatment with polyphenols reduces both acute and sensitized locomotor response to high dose morphine, with minimal effects on low dose sensitization (**Fig. 1**). On morphine CPP, BDPP pretreatment markedly reduced formation of preference at low dose morphine, but potentiated it at high doses (**Fig. 2**). When we examined the NAc, we found transcriptional effects of morphine were potentiated by BDPP treatment particularly at the high dose of morphine (**Fig. 3**). Detailed examination of gut microbiome composition and function reinforced significant morphine dose by polyphenol interactions, with robust effects of polyphenol treatment seen after low but not high dose morphine (**Fig. 4**). Multiple mechanistic studies consequently identified that these myriad effects were not simply explained by SIRT1 activation or by a subset of BDPP derived metabolites (**Fig. 5**). Taken together, we find polyphenol treatment has both marked and complex effects on the behavioral and physiological response to morphine. These studies lay foundations for future translational work examining the potential for phytochemicals as novel treatments against opioid use disorders.

Based on review of the literature, these studies are the first to examine the effects of polyphenols on addiciton-like behaviors in an opioid model. Multiple studies demonstrate that polyphenols and particularly resveratrol can reduce the development of tolerance for morphine, and have anti-nociceptive effects in models of pain [26–28,53–56]. Additionally, oral consumption of BDPP was previously reported to attenuate pain associated with lumbar interverterbral disc injury in rats [24]. While our experiments did not address issues of tolerance or analgesia from morphine, it is possible that the underlying mechanisms are similar, and future work in this area can assess overlap in these areas.

Several studies have examined the effects of polyphenols on models of stimulant use disorders with conflicting results. A study examining the effects of SIRT1 on behavioral responses to cocaine found resveratrol enhances cocaine CPP via a SIRT1 dependent mechanism [57]. However, this effect was tested only at a single (5mg/kg) dose of cocaine, and the resveratrol treatment differed markedly from the one used in our studies. Conversley, resveratrol was also reported to reduce the formation of CPP for high dose cocaine (15mg/kg) [58]. These seemingly opposing effects on cocaine related behaviors lend further evidence to complex drug dose x polyphenol effects on addiction-like behaviours as observed in the current study (**Figs. 1-2**). Such findings indicate the effects of polyphenols on addiction-related behaviours are likley influenced by factors including the drug of abuse in question, dose, chronic or acute exposure, polyphenol used and route of administration.

There is a wealth of literature demonstrating that prolonged changes in gene expression in key limbic brain structures underlie behavioral and synaptic plasticity in models of opioid and other substance use disorders [7,8]. Here, we find that treatment with the BDPP alters gene expression patterns in the NAc, paricularly in response to high dose morphine (**Fig. 3**). The exact mechanism of the gene expression changes we see following BDPP and high dose morphine are not clear. Previous studies have reported gene expression changes in the brain following polyphenol treatment, focusing on potential epigenetic effects of polyphenols [59], including regulation of DNMTs and histone deacetylases [59–61].

In addition to these behavioral and transcriptomic effects, we found effects of BDPP treatment on the contents and function of the gut microbiome – most notably at low dose morphine (**Fig. 4**). It is established that dietary polyphenols carry prebiotic properties, enhancing the growth of specific bacterial species which elicit health benefits to the host [49,50]. Indeed, we noted increased abundance of beneficial bacteria including *Akkermansia*, as a result of BDPP treatment in low dose morphine groups (**Fig. 4H**). BDPP induced alterations to the microbiome represent a potential mechanism for the BDPP behavioral responses we report.

In summary, we demonstrate treatment with dietary polyphenols markedly alters behavioral response to opioids with strong dose x treatment interactions. Dose and treatment effects were also seen on transcriptomic analyiss of the NAc and microbiome sequencing. Mechanistic studies were unable to simplify these effects to SIRT1 activation or a subset of BDPP metabolites. However, importantly, our data demonstrate the capability of dietry polyphenols to modulate behavioural and molecular responses to morphine in a model of opioid use disorder. There remains an urgent need to develop new interventions for treatment of OUD, and in particular non-traditional targets lacking intrinsic abuse potential. The data presented herein lay a strong foundation for the importance of dietary polyphenols as a potential translational research target in OUD, and future studies aim to dissect mechanism, dose, and timing interactions to best harness the beneficial properties of dietary phytochemicals.

## Acknowledgements

Morphine was provided by NIDA drug supply. Schematics in all figures were prepared using BioRender.com with full permission to publish.

## Author Contributions

D.D.K and A.O. designed the experiments. A.O., R.S.H., K.R.M., Y.A.D., S.M.Z., A.L.S., and K.E.L. performed experiments. A.O. & D.D.K analyzed data. A.O., Y.A.D., S.M.Z., A.L.S., K.E.L., and D.D.K designed and created figures. K.J.T., T.O., and G.M.P. contributed critical reagents and assisted in assay design. D.D.K and A.O. wrote the paper. All authors provided critical edits on the final draft of the manuscript.

## Supplementary Information

### Supplementary Methods

#### Locomotor Sensitization

The locomotor arena consisted of a frame crossed with infrared beams in the x and y dimensions. Clean, empty rat cages were placed within this frame to contain the mice while simultaneously allowing penetration of the infrared beams. Animals could freely move throughout the space and infrared beam breaks were counted to determine locomotor activity. Control or BDPP treated mice were injected with saline, 5 mg/kg, or 15 mg/kg morphine and their activity monitored for the subsequent 45 mins. For “challenge injection” experiments mice were tested for five days before being returned to their home cages for ten days of abstinence. Ten days later they were given a morphine challenge injection of the same dose as the initial injections and were placed back into the locomotor arena to measure persistence of morphine locomotor sensitization.

#### Morphine conditioned place preference

Each CPP apparatus consists of 3 distinct chambers: a middle, small entry chamber and two larger conditioning chambers on either side of the entry. Conditioning chambers were distinct in wall color and floor texture. The left end chamber had gray walls and a large grid floor, and the right end chamber had black and white striped walls and a small grid floor. Conditioned place preference occurred over 5 days: day 1 was a pre-test, days 2-4 were conditioning days, and the fifth day was the post test. On pre-test day, mice were placed into the center chamber of the apparatus and were allowed to explore all 3 chambers for 20 mins. Time spent in each chamber was recorded; mice spending > 70 % of their time in one chamber were excluded. Mice were assigned to their morphine-paired chamber using an unbiased approach such that group preference on pre-test was as close to zero as possible. On conditioning days, mice were injected with saline subcutaneously in the morning and confined to one end chamber and injected with either 2.5, 5, or 15 mg/kg morphine and confined in the opposite end chamber in the afternoon. Both morning and afternoon conditioning sessions lasted 45 mins. On test day, mice were again allowed to explore all 3 chambers of the apparatus for 20 mins, as described above for pre-test. Place preference score was calculated as: time spent in the morphine chamber on day 5 – time spent day 1 (Test – PreTest).

#### RNA Sequencing

RNA from frozen NAc punches was isolated using RNeasy kits (Qiagen-#74106) with on-column DNAase digestion (#79254) per manufacturer’s protocol. The integrity and purity of total RNA were assessed using Agilent Bioanalyzer and OD260/280 using Nanodrop. The RNA sequencing library was generated using NEBNext Ultra II RNA library Prep Kit for Illumina using manufacturer’s instructions (New England Biolabs, Ipswich, MA, USA). The sequencing library was validated on the Agilent TapeStation (Agilent Technologies, Palo Alto, CA, USA), and quantified by using Qubit 2.0 Fluorometer (ThermoFisher Scientific, Waltham, MA, USA) as well as by quantitative PCR (KAPA Biosystems, Wilmington, MA, USA). The libraries were then multiplexed and sequenced on an Illumina HiSeq 4000 instrument using a 2×150bp Paired End (PE) configuration according to manufacturer’s instructions. Image analysis and base calling were conducted by the HiSeq Control Software (HCS). Raw sequence data (.bcl files) generated from Illumina HiSeq was converted into fastq files and de-multiplexed using Illumina bcl2fastq 2.20 software. One mismatch was allowed for index sequence identification.

#### RNA-sequencing Data Analysis

The raw RNA-Seq reads (Fastq files) for each sample were aligned and read counts generated using the cloud-based BioJupies software package using the default settings [1]. Raw count matrices were then analyzed using the Network Analyst Software package with the variance filter set to 5% and the low abundance filter set to 2 [2,3]. Data were normalized to Log2 counts per million. Differential gene expression analysis was then performed using Deseq2. Statistical significance was set at a threshold of FDR-corrected *p* < 0.2 except as described. For pathway and transcription factor analysis all genes marked as “predicted gene”, “pseudogene”, Riken gene, and ribosomal protein encoding genes were removed prior to analysis. Identification of significantly enriched gene ontologies was performed using the g:Profiler analysis package [4]. For identification of enrichment of predicted transcription factor activity we utilized the Enrichr software package querying the Chea and ENCODE datasets to identify targets based on publicly available ChIP-seq datasets [5,6]. Volcano plots were made using GraphPad Prism.

#### 16s-Sequencing of Cecal Content

Bacterial genomic DNA was isolated from frozen cecal samples using the DNeasy PowerSoil Pro kit (Qiagen) according to standard protocols with a modification that included an extended bead beating step. Extracted DNAs were checked for quality and quantity by spectrophotometric measurements with NanoDrop (ThermoFisher Scientific Inc). PCR amplification was subsequently achieved using primers 341F (5’-CCTACGGGNGGCWGCAG-3’) and 805R (5’-GACTACHVGGGTATCTAATCC-3’) targeting regions (V3-V4) of the 16S rRNA gene. The libraries were sequenced using illumina NovaSeq (2 × 250 bp paired-end) platform using standard parameters.

#### 16s-Sequencing Data Analysis

Amplicons were trimmed, merged using FLASH [7] and chimera filtered using Vsearch (v2.3.4). Sequences with ≥97% similarity were assigned to the same operational taxonomic units (OTUs). Representative sequences were chosen for each OTU, followed by taxonomic assignment using the RDP (Ribosomal Database Project) classifier. The differences of the dominant species in different groups and multiple sequence alignment were conducted by mafft software (v7.310). OTU abundance information was used to determine alpha diversity as the Chao1 metric. Principle coordinates analysis plots were generated using the Unifrac distance as an assessment of beta diversity, both using the Quantitative Insights Into Microbial Ecology (QIIME) package v1.8.0 [8]. Statistical significance for alpha diversity was analyzed using a 2×2 between subjects ANOVA with morphine dose and drink solution provided as fixed factors. At the phylum and genus level, any bacterial groups with mean expression levels ≥ 0.05% abundance and a Wilcoxon’s FDR corrected *p* value < 0.2 were considered statistically significant.

#### Microbial Functional Profile Prediction and LEfSe methods

Predicted functional profiles of bacterial groups were determined using Phylogenetic Investigation of Communities by Reconstruction of Unobserved States (PICRUSt2) bioinformatics software package as published [9]. Firstly, the OTU table produced within QIIME was normalized for multiple 16s RNA copy numbers. Then the obtained normalized OTU table was placed into a reference tree in order to obtain the Kyoto Encylopedia of Genes and Genomes (KEGG) orthologs (KO). In order to assess the functional pathways, including enzymatic pathways affected by microbiome changes due to BDPP treatment and morphine dose, sequence counts of all KO pathways for each sample were converted to mean proportion percent using the formula: (sequence count for specific KO pathway / total sequence count of all pathways x 100). Subsequently a two-tailed students *t*-test and FDR < 0.01 was applied to all pathways. K identifiers of KO pathways which met this significance criteria were then uploaded into the KEGG database www.genome.jp/kegg/ko.html to identify mapped cellular functions.

OTU tables generated were further subjected to Linear discriminant analysis (LDA) effect size (LEfSe) to calculate the taxa that best discriminated between control and BDPP treatment groups. Specifically, the non-parametric factorial Kruskal-Wallis (KW) sum-rank test was used to detect taxa with significant differential abundance among groups. Taxa consistency was then investigated using a set of pairwise tests among subclasses with the (unpaired) Wilcoxon rank-sum test. Finally, LEfSe uses LDA to estimate the effect size of each differentially abundant taxa. Taxa that reached a linear discriminant analysis score (log10) >3.0 and p < 0.05 are reported. Taxa that discriminating between treatment groups are visualized on taxonomic trees called Cladograms. Cladograms show how the taxa that discriminate between the various treatment groups are related to each other in terms of taxonomy and ontologies of functional pathways.

### Supplementary Results

#### Acute and Sensitized Locomotor Responses

Given the reduced response of the BDPP treated group to acute 15mg/kg morphine, we analyzed this response with increased granularity across this session. When the locomotor response is examined in five minute bins across the session, the main effect of treatment immediately becomes clear between the two morphine groups (F_(1,10)_=9.98; *p*=0.01), with post-hoc testing showing that BDPP-treated mice had decreased locomotor activity compared to controls starting in the second bin (**Fig. S1A**). There was no main effect of time across the session (F_(8,80)_=1.28; *p*=0.26) or time x treatment interaction (F_(8,80)_=1.11; *p*=0.37) for the two morphine groups – the saline groups are provided for visual comparison in this panel but not directly compared to morphine groups.

As locomotor sensitization has been shown to persist for weeks, we next performed an experiment in which previously injected mice were given a single challenge injection of morphine two weeks after the conclusion of the initial sensitization experiment. In this context we find that there is no main effect of BDPP treatment (**Fig. S1B** - F_(1,35)_=0.07; *p*=0.80) or significant treatment x dose interaction (F_(2,35)_=0.08; *p*=0.92). As expected, there was a robust effect of morphine dose (F_(2,35)_=21.77; *p*<0.0001). Taken together, these results suggest that dietary polyphenols have marked effects on locomotor activation in response to morphine, with notable dose and timing interactions.

#### Conditioned place preference following withdrawal

Since previous studies have demonstrated that conditioned place preference can be potentiated in animals that have previously been treated with morphine [10], we performed CPP after morphine withdrawal as described in **Fig. 2**. However, to ensure that we saw a main effect of morphine withdrawal, we compared effects of CPP completed in previously naïve mice, to those who underwent morphine pretreatment and withdrawal. Here we find that there is indeed a main effect of withdrawal (**Fig. S2** – F_(1,69)_=4.48; *p* = 0.03), as well as BDPP treatment (F_(1,69)_=12.57; *p* = 0.0007), with no significant interactions.

#### Predicted Functional Enzymatic Pathways

Given the distinct effects of BDPP treatment on microbiome composition in the low and high dose chronic morphine groups, analysis of functional enzymatic pathways differentially affected by BDPP treatment and morphine dose was conducted. Direct comparison of mean sequence proportions between BDPP-5mg/kg and control 5mg/kg morphine groups revealed 222 significantly different pathways (two-tailed Student’s t-test and FDR corrected *p* <0.01). However, direct comparison of mean sequence proportions between BDPP-15mg/kg and control 15mg/kg morphine groups revealed no significantly different pathways using the same cut off thresholds (Full PICRUSt2 list is supplied in **Table S12)**. In order to identify which cellular functions the significant enzymes in the 5mg/kg morphine group mapped to, KEGG database was utilized to annotate the significant pathways identified. Interestingly a significant decrease in the mean proportion of enzymes involved in Glycolysis/gluconeogenesis were observed in BDPP 5mg/kg morphine treated mice **(Fig. S3C)**. Meanwhile a significant increase in a number of enzymes involved in Fatty Acid Metabolism and Amino Acid Metabolism was also observed in BDPP 5mg/kg morphine treated mice.

#### Correlations between CPP and 16s results

As a marker of which bacterial groups may be playing a role in BDPP mediated behavioral responses to high and low dose morphine, we performed correlation analysis between caecal abundance of select bacterial phyla and genus (significantly changed by treatment), with preference in the CPP paradigm. Correlation matrices with exact r and p values for the select phyla and genus correlations are available in supplementary data **Table S14** and **S15** respectively. A heatmap of Spearman’s r values for phyla and genus are presented in **Fig. S4A** and **S5A** respectively. Interestingly, at the phylum level Bacteroidetes showed a significant negative linear correlation with CPP across all treatment groups (**Fig. S4B** Spearman *r* = -0.043, *p* = 0.01). Indicating a higher abundance of Bacteroidetes may be protective of morphine CPP. When we correlated individual treatment groups separately, Bacteroidetes showed a significant negative correlation only in control treated groups (**Fig. S4C** Spearman *r* = -0.691, *p* = 0.004). While the abundance of Firmicutes did not show a significant correlation with CPP across all treatment groups, there was a strong trend towards a positive linear correlation (**Fig. S4D** Spearman *r* = -0.34, *p* = 0.05). Suggesting a higher abundance of Firmicutes contributes to the development of morphine CPP. Correlation of Firmicutes with CPP score within individual treatment groups showed a significant positive correlation only between control treated animals (**Fig. S4C** Spearman *r* =0.595, *p* = 0.016). Furthermore, the phyla Proteobacteria showed a significant negative linear correlation with CPP across all treatment groups (**Fig. S4F** Spearman *r* = -0.492, *p* = 0.00). Within individual treatment groups, Proteobacteria showed a significant negative correlation between 15mg/kg morphine treated animals (**Fig. S4G** Spearman *r* =-0.652, *p* = 0.08) and control treated groups (**Fig. S4H** Spearman *r* =-0.708, *p* = 0.03).

At the genus level, abundance of *Muribaculaceae* was found to significantly negatively correlate with CPP score when comparing all treatment groups (**Fig. S5B** Spearman *r* = -0.431, *p* = 0.014). When we correlated individual treatment groups separately at the genus level, only control treated animals showed a significant negative correlation between *Muribaculaceae* and CPP (**Fig. S5C** Spearman *r* = -0.759, *p* = 0.001. Similarly, abundance of *Parasutterella* was also found to negatively correlate with CPP score when comparing all treatment groups (**Fig. S5D** Spearman *r* = -0.433, *p* = 0.013). Within individual treatment groups, *Parasutterella* showed a significant negative correlation between 15mg/kg (**Fig. S5E** Spearman *r* = -0.5575, *p* = 0.02) and control treated animals (**Fig. S5F** Spearman *r* = -0.6992, *p* = 0.003).

These correlation results are in line with findings from LEfSE analysis. *LEfSe* showed the microbiome of animals which develop morphine CPP, namely the 5mg/kg control and BDPP 15mg/kg, is characterized by a Firmicutes dominant enterotype (**Fig. 4I** – **green** and **Fig. 4J-red**). Here we see Firmicutes abundance show a strong trend towards a positive correlation with CPP. While the microbiome of animals which do not develop morphine CPP, namely BDPP 5mg/kg and control 15mg/kg, is characterized by a Proteobacteria/*Parasutterella* and Bacteroidetes/*Muribaculaceae* enterotype (**Fig. 4I** – **red** and **Fig. 4J-green**). Here we show these bacterial taxa negatively correlate with CPP. Taken together, 16s analysis suggest alterations to key bacterial clusters, namely Bacteroidetes/*Muribaculaceae*, Proteobacteria/ *Parasutterella* and Firmicutes/*Clostridium* may play a role in mediating the effect of BDPP treatment on behavioural responses to high and low dose morphine.

### Supplemental Discussion

In addition to models of substance use disorders, polyphenols have also been shown to have marked protective effects in multiple models of neuropsychiatric disorders[11,12]. For example, in models of Alzhimers disease (AD) polyphenolic compounds from a variety of diverse sources have the ability to improve cognitive function and reduce neuropathology in animal models of AD through multiple mechanismsm including improving synaptic plasticity [13]. In particular, a combination of resveratrol, GSPE and concord grape juice have been shown to synergistically mitigate Amyloid β peptide -mediated neuropathology and cognitive impairments in the brain in a mouse model of AD [14]. In addition to effects on neurodegenerative diseases, there have also been a number of studies showing effects of polyphenols on models of depression. Specifically, chronic treatment with BDPP was found to protect against stress induced depression in mice via an upregulation of VGF in the hippocampus [15], similar to more traditional antidepressants[16]. Chronic BDPP treatment was also reported to alleviate chronic social defeat stress-induced social avoidance via two BDPP-derived metabolites, Mal-Gluc and DHCA, which were identified to promote resilience by modulating synaptic plasticity and peripheral inflammation [17]. Taken together, this body of literature describes important effects of phytochemicals in modulating neuroplasticity in a myriad of neuropsychiatric conditions.

Regarding effects on the microbiome and its functional output, dietary polyphenols have been shown to promote generation of short chain fatty acids (SCFAs) [18]. These byproducts of bacterial fermentation of fibers have numerous effects on brain and behavior [19]. Importantly, previous work from our lab has shown that the presence of SCFA’s is critical for behavioral responses to both cocaine and opioids [20,21]. It is possible that behavioral effects of BDPP treatment could be caused by polyphenols’ production of key microbial metabolites such as the SCFAs. Indeed, predictive functional analysis of 16s data revealed a significant increase in enzymes involved in SCFA metabolism in the low dose morphine BDPP treated group compared to control counter parts (Fig.**S3C**). Additionally, polyphenols are known to possess antibacterial effects and have been shown to decrease the Firmicutes/Bacteroidetes ratio [22]. This is in line with the results from the current study showing a decrease in Firmicutes/Bacteroidetes ratio following BDPP treatment at low dose morphine (**Fig. 4G**).

## Supplementary Figures

**Figure S1.**
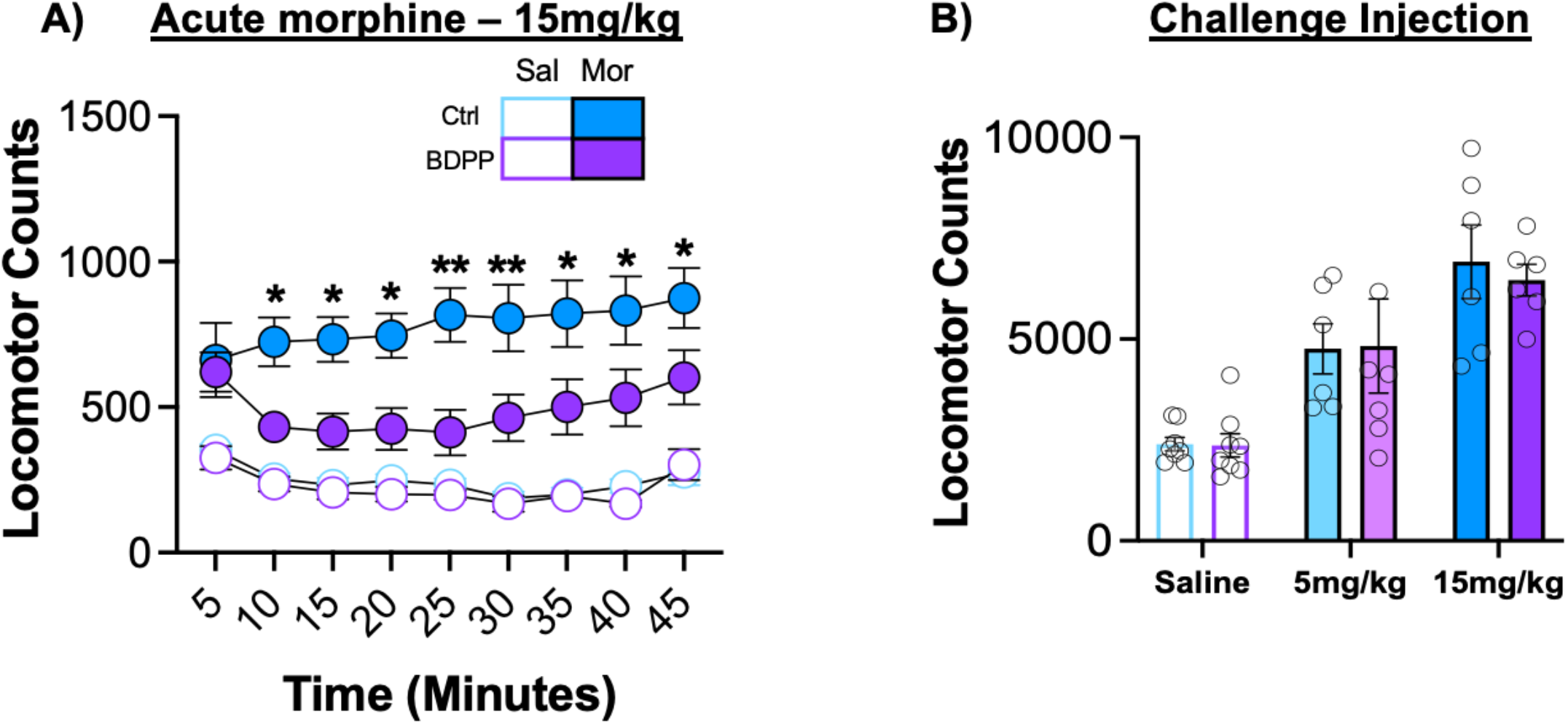
Acute and Sensitized Locomotor Responses. **(A)** Acute effects of morphine on locomotor activation were also decreased by pre-treatment with BDPP polyphenols. **(B)** In mice previously treated with 5 days of morphine, polyphenol treatment did not affect persistence of morphine sensitization after two weeks of withdrawal.

**Figure S2.**
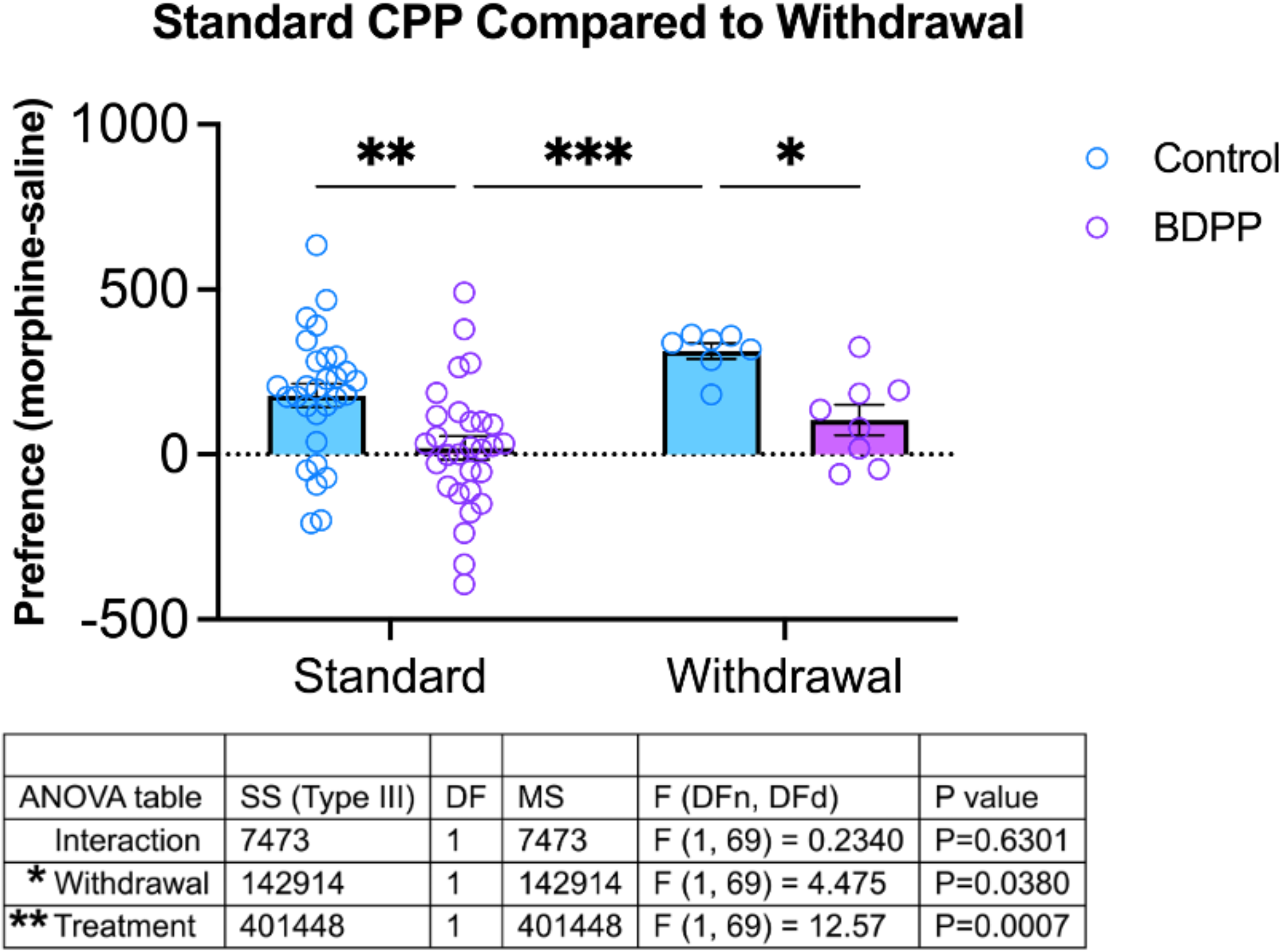
Effect of morphine pretreatment on subsequent development of conditioned place preference. Mice that were pretreated with morphine followed by two weeks of withdrawal prior to CPP conditioning demonstrated more robust acquisition of conditioned place preference on subsequent training.

**Figure S3.**
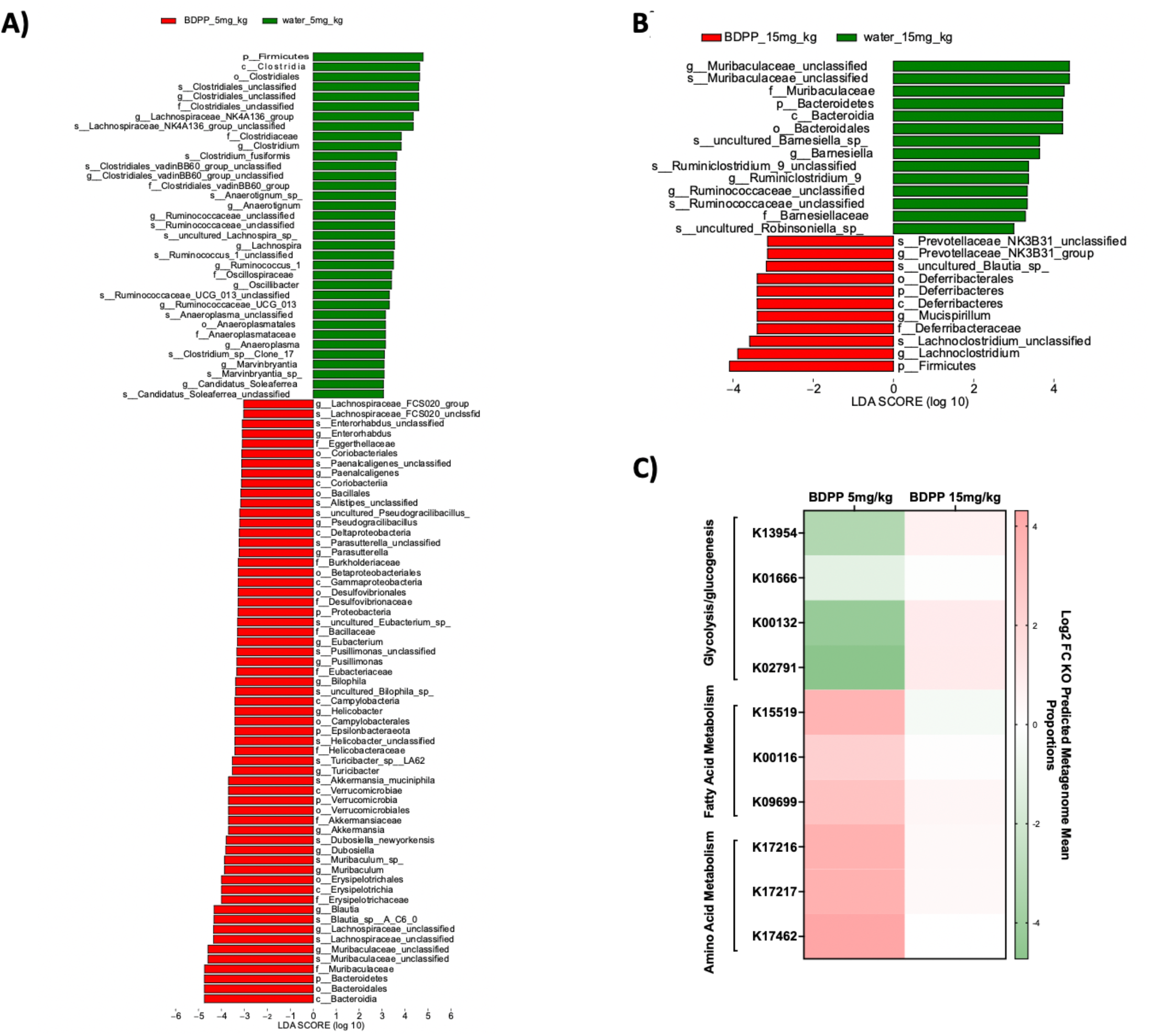
BDPP Treatment Differentially Regulates Microbiome Composition and Functional Output in Low and High Dose Morphine Treated Animals. Linear discriminant analysis (LDA) effect size (LEfSe) analysis of 16S data was used to determine bacterial taxa which are differentially represented between treatment groups **(A)** list of bacterial taxa found to be overrepresented (Red) or underrepresented (Green) in BDPP 5gm/kg relative to control 5mg/kg treatment groups **(B)** list of bacterial taxa found to be overrepresented (Red) or underrepresented (Green) in BDPP 15mg/kg relative to control 15mg/kg treatment groups **(C)** Phylogenetic Investigation of Communities by Reconstruction of Unobserved States (PICRUSt2) package identified enzymes predicted to be involved in glycolysis/glucogenesis to be significantly decreased, fatty acid metabolism and amino acid metabolism enzymes increased in BDPP 5mg/kg treated animals relative to controls (FDR corrected *p* <0.01 significance cut off).These same enzymatic pathways were unchanged in BDPP 15mg/kg treated animals.

**Figure S4.**
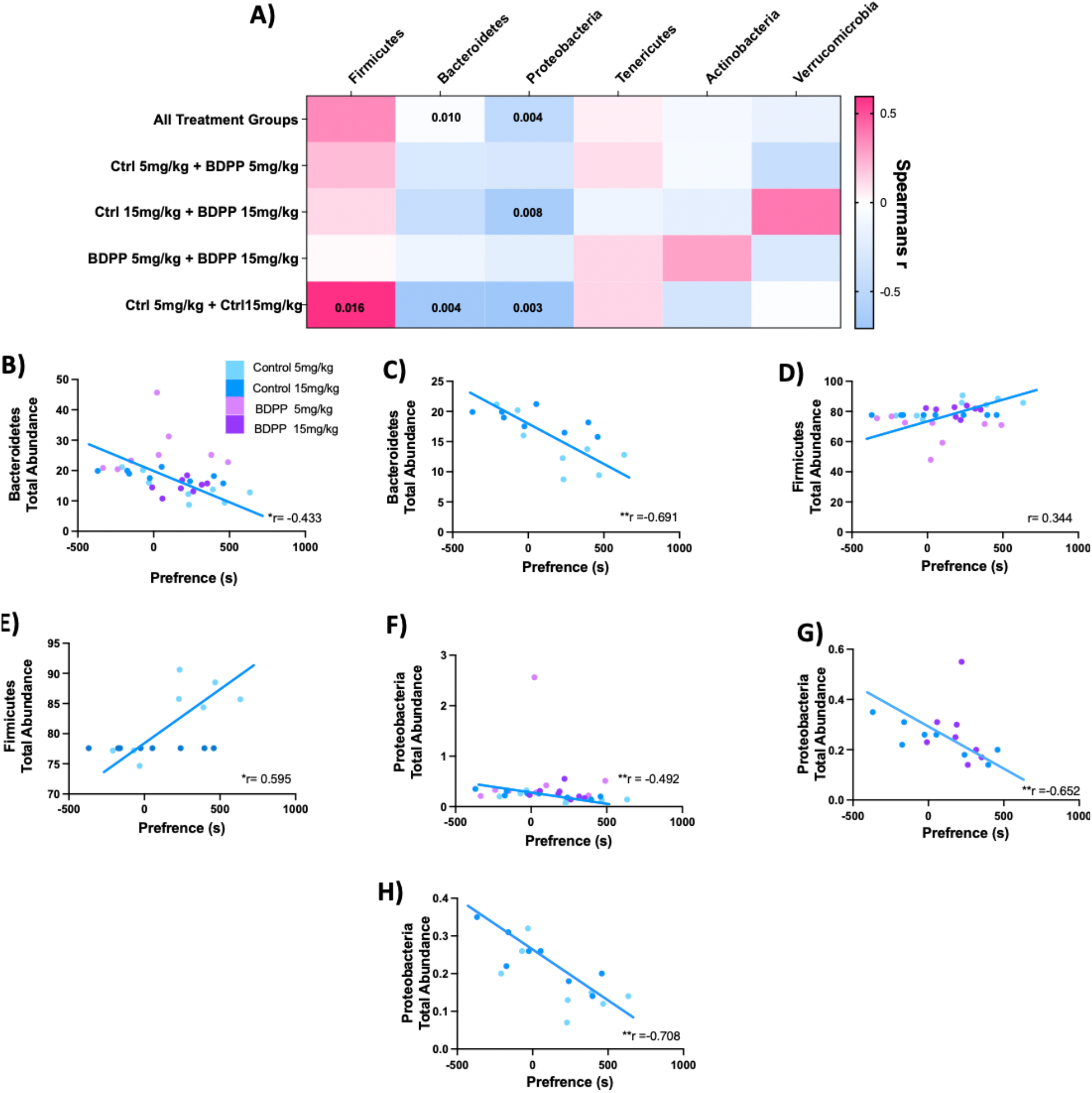
Correlations between Phyla Total Abundance and CPP Score. Mice that were pre-treated with BDPP for two weeks followed by CPP testing had cecal content collected 24 hours after test day for 16s analysis **(A)** Correlation heatmap of select phyla (columns) with CPP preference, across all treatment groups or within individual treatment groups (rows). Exact r values for each phylum and exact p values are available in **Supplementary Table S14. (B)** Bacteroidetes correlate with CPP across all treatment groups (Spearman *r* = -0.04, *p* = 0.01) and **(C)** correlate in control 5mg/kg vs control 15mg/kg (Spearman *r* = -0.69, *p* = 0.04) **(D)** Firmicutes show a strong trend towards a significant correlation with CPP across all treatment groups (Spearman *r* = 0.34, *p* = 0.05) and **(E)** significantly correlate in control 5mg/kg vs control 15mg/kg (Spearman *r* = 0.60, *p* = 0.02). **(F)** Proteobacteria correlate with CPP across all treatment groups (Spearman *r* = -0.49, *p* = 0.00) and **(G)** correlate in control 15mg v BDPP 15mg (Spearman *r* = -0.65, *p* = 0.01) and **(H)** correlate in control 5mg/kg v control 15mg/kg (Spearman *r* = -0.71, *p* = 0.00).

**Figure S5.**
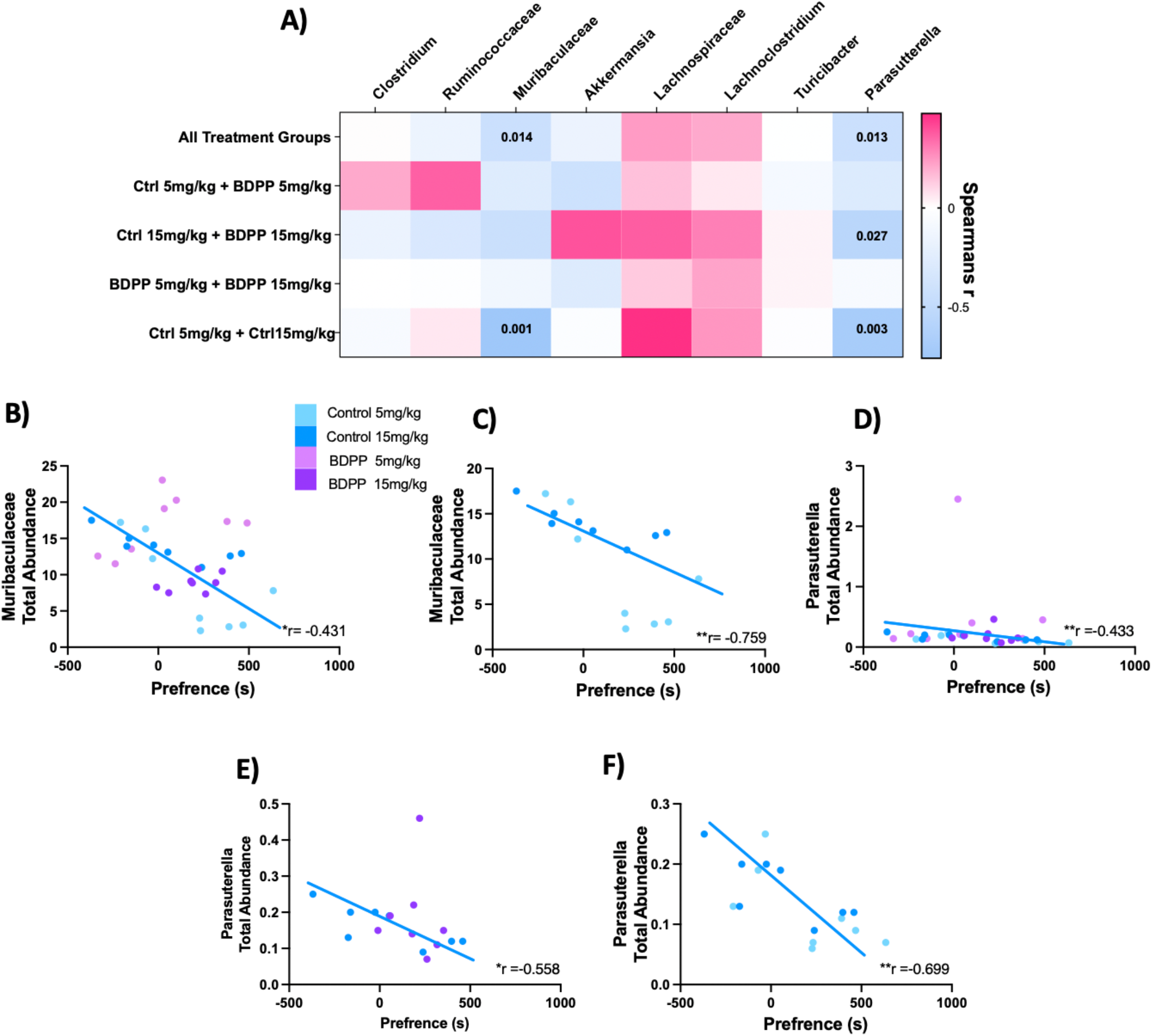
Correlations between Genus Total Abundance and CPP Score. Mice that were pre-treated with BDPP for two weeks followed by CPP testing had caecal content collected 24 hours after test day and sent for 16s analysis **(A)** Correlation heatmap of individual select genus (columns) with CPP preference across all treatment groups or within individual treatment groups (rows). Exact r values for each genus and exact p values are available in Supplementary Table **S15. (B)** *Muribaculaceae* correlate with CPP across all treatment groups (Spearman *r* = -0.43, *p* = 0.01) and **(C)** correlate in control 5mg/kg vs control 15mg/kg (Spearman *r* = -0.76, *p* = 0.001) **(D)** *Parasutterella* correlate with CPP across all treatment groups (Spearman *r* = - 0.43, *p* = 0.01) and **(E)** correlate in control 15mg v BDPP 15mg (Spearman *r* = - 0.56, *p* = 0.03) and **(F)** correlate in control 5mg/kg v control 15mg/kg (Spearman *r* = -0.7, *p* = 0.003).

## References

1. Koob GF, Volkow ND. Neurocircuitry of Addiction. Neuropsychopharmacol. 2010;35:217–238.

2. Le Moal M, Koob GF. Drug addiction: Pathways to the disease and pathophysiological perspectives. European Neuropsychopharmacology. 2007;17:377–393.

3. Hedegaard H, Brigham Bastian, Trinidad J, Spencer M, Warner M. Drugs Most Frequently Involved in Drug Overdose Deaths: United States, 2011–2016. National Vital Statistics Reports. 2018;67:14.

4. Products - Vital Statistics Rapid Release - Provisional Drug Overdose Data. 2022. https://www.cdc.gov/nchs/nvss/vsrr/drug-overdose-data.htm. Accessed 13 May 2022.

5. Manchikanti L, Vanaparthy R, Atluri S, Sachdeva H, Kaye AD, Hirsch JA. COVID-19 and the Opioid Epidemic: Two Public Health Emergencies That Intersect With Chronic Pain. Pain Ther. 2021;10:269–286.

6. Schuckit MA. Treatment of Opioid-Use Disorders. New England Journal of Medicine. 2016;375:357–368.

7. Browne CJ, Godino A, Salery M, Nestler EJ. Epigenetic Mechanisms of Opioid Addiction. Biological Psychiatry. 2020;87:22–33.

8. Robison AJ, Nestler EJ. Transcriptional and epigenetic mechanisms of addiction. Nat Rev Neurosci. 2011;12:623–637.

9. Wolf ME. Synaptic mechanisms underlying persistent cocaine craving. Nat Rev Neurosci. 2016;17:351– 365.

10. Lacagnina MJ, Rivera PD, Bilbo SD. Glial and Neuroimmune Mechanisms as Critical Modulators of Drug Use and Abuse. Neuropsychopharmacology. 2017;42:156–177.

11. O’Sullivan SJ, Malahias E, Park J, Srivastava A, Reyes BAS, Gorky J, et al. Single-Cell Glia and Neuron Gene Expression in the Central Amygdala in Opioid Withdrawal Suggests Inflammation With Correlated Gut Dysbiosis. Frontiers in Neuroscience. 2019;13.

12. Wang J, Tang C, Ferruzzi MG, Gong B, Song BJ, Janle EM, et al. Role of standardized grape polyphenol preparation as a novel treatment to improve synaptic plasticity through attenuation of features of metabolic syndrome in a mouse model. Molecular Nutrition & Food Research. 2013;57:2091–2102.

13. Ward L, Pasinetti GM. Recommendations for Development of Botanical Polyphenols as “Natural Drugs” for Promotion of Resilience Against Stress-Induced Depression and Cognitive Impairment. Neuromol Med. 2016;18:487–495.

14. Zhao W, Wang J, Bi W, Ferruzzi M, Yemul S, Freire D, et al. Novel application of brain-targeting polyphenol compounds in sleep deprivation-induced cognitive dysfunction. Neurochemistry International. 2015;89:191–197.

15. Shukitt-Hale B, Carey A, Simon L, Mark DA, Joseph JA. Effects of Concord grape juice on cognitive and motor deficits in aging. Nutrition. 2006;22:295–302.

16. Wang J, Hodes GE, Zhang H, Zhang S, Zhao W, Golden SA, et al. Epigenetic modulation of inflammation and synaptic plasticity promotes resilience against stress in mice. Nat Commun. 2018;9:477.

17. Wang J, Ho L, Zhao W, Ono K, Rosensweig C, Chen L, et al. Grape-derived polyphenolics prevent Abeta oligomerization and attenuate cognitive deterioration in a mouse model of Alzheimer’s disease. J Neurosci. 2008;28:6388–6392.

18. Bahi A, Nurulain SM, Ojha S. Ethanol intake and ethanol-conditioned place preference are reduced in mice treated with the bioflavonoid agent naringin. Alcohol. 2014;48:677–685.

19. Yunusoğlu O. Evaluation of the effects of quercetin on the rewarding property of ethanol in mice. Neurosci Lett. 2022;768:136383.

20. Yunusoğlu O. Resveratrol impairs acquisition, reinstatement and precipitates extinction of alcohol-induced place preference in mice. Neurol Res. 2021;43:985–994.

21. Chen H, Wu J, Zhang J, Fujita Y, Ishima T, Iyo M, et al. Protective effects of the antioxidant sulforaphane on behavioral changes and neurotoxicity in mice after the administration of methamphetamine. Psychopharmacology. 2012;222:37–45.

22. Zeng Q, Xiong Q, Zhou M, Tian X, Yue K, Li Y, et al. Resveratrol attenuates methamphetamine-induced memory impairment via inhibition of oxidative stress and apoptosis in mice. J Food Biochem. 2021;45:e13622.

23. Sun D, Yue Q, Guo W, Li T, Zhang J, Li G, et al. Neuroprotection of resveratrol against neurotoxicity induced by methamphetamine in mouse mesencephalic dopaminergic neurons. Biofactors. 2015;41:252– 260.

24. Lai A, Ho L, Evashwick-Rogler TW, Watanabe H, Salandra J, Winkelstein BA, et al. Dietary polyphenols as a safe and novel intervention for modulating pain associated with intervertebral disc degeneration in an in-vivo rat model. PLoS One. 2019;14:e0223435.

25. Ma Y, Liu S, Shu H, Crawford J, Xing Y, Tao F. Resveratrol alleviates temporomandibular joint inflammatory pain by recovering disturbed gut microbiota. Brain Behav Immun. 2020;87:455–464.

26. Hirata K, Nishiki Y, Goto R, Inagaki M, Oshima K, Shimazu Y, et al. Resveratrol suppresses nociceptive jaw-opening reflex via 5HT3 receptor-mediated GABAergic inhibition. Neurosci Res. 2020;160:25–31.

27. Iqubal A, Ahmed M, Iqubal MK, Pottoo FH, Haque SE. Polyphenols as Potential Therapeutics for Pain and Inflammation in Spinal Cord Injury. Curr Mol Pharmacol. 2021;14:714–730.

28. Gupta YK, Sharma M, Briyal S. Antinociceptive effect of trans-resveratrol in rats: Involvement of an opioidergic mechanism. Methods Find Exp Clin Pharmacol. 2004;26:667–672.

29. Pasinetti GM, Wang J, Marambaud P, Ferruzzi M, Gregor P, Knable LA, et al. Neuroprotective and metabolic effects of resveratrol: therapeutic implications for Huntington’s disease and other neurodegenerative disorders. Exp Neurol. 2011;232:1–6.

30. Jang I-A, Kim EN, Lim JH, Kim MY, Ban TH, Yoon HE, et al. Effects of Resveratrol on the Renin-Angiotensin System in the Aging Kidney. Nutrients. 2018;10:1741.

31. Ayissi VBO, Ebrahimi A, Schluesenner H. Epigenetic effects of natural polyphenols: A focus on SIRT1-mediated mechanisms. Molecular Nutrition & Food Research. 2014;58:22–32.

32. Frolinger T, Herman F, Sharma A, Sims S, Wang J, Pasinetti GM. Epigenetic modifications by polyphenolic compounds alter gene expression in the hippocampus. Biology Open. 2018;7:bio035196.

33. Jiang C, Sakakibara E, Lin W-J, Wang J, Pasinetti GM, Salton SR. Grape-derived polyphenols produce antidepressant effects via VGF- and BDNF-dependent mechanisms. Annals of the New York Academy of Sciences. 2019;1455:196–205.

34. Dasgupta B, Milbrandt J. Resveratrol stimulates AMP kinase activity in neurons. Proceedings of the National Academy of Sciences. 2007;104:7217–7222.

35. Pasinetti GM, Singh R, Westfall S, Herman F, Faith J, Ho L. The Role of the Gut Microbiota in the Metabolism of Polyphenols as Characterized by Gnotobiotic Mice. J Alzheimers Dis. 2018;63:409–421.

36. Hofford RS, Mervosh NL, Euston TJ, Meckel KR, Orr AT, Kiraly DD. Alterations in microbiome composition and metabolic byproducts drive behavioral and transcriptional responses to morphine. Neuropsychopharmacology. 2021. 14 June 2021. https://doi.org/10.1038/s41386-021-01043-0.

37. Raudvere U, Kolberg L, Kuzmin I, Arak T, Adler P, Peterson H, et al. g:Profiler: a web server for functional enrichment analysis and conversions of gene lists (2019 update). Nucleic Acids Res. 2019;47:W191–W198.

38. Chen EY, Tan CM, Kou Y, Duan Q, Wang Z, Meirelles GV, et al. Enrichr: interactive and collaborative HTML5 gene list enrichment analysis tool. BMC Bioinformatics. 2013;14:128.

39. Kuleshov MV, Jones MR, Rouillard AD, Fernandez NF, Duan Q, Wang Z, et al. Enrichr: a comprehensive gene set enrichment analysis web server 2016 update. Nucleic Acids Research. 2016;44:W90–W97.

40. Caporaso JG, Kuczynski J, Stombaugh J, Bittinger K, Bushman FD, Costello EK, et al. QIIME allows analysis of high-throughput community sequencing data. Nat Methods. 2010;7:335–336.

41. Douglas GM, Maffei VJ, Zaneveld JR, Yurgel SN, Brown JR, Taylor CM, et al. PICRUSt2 for prediction of metagenome functions. Nat Biotechnol. 2020;38:685–688.

42. Simpson GR, Riley AL. Morphine preexposure facilitates morphine place preference and attenuates morphine taste aversion. Pharmacology Biochemistry and Behavior. 2005;80:471–479.

43. Blaze J, Wang J, Ho L, Mendelev N, Haghighi F, Pasinetti GM. Polyphenolic Compounds Alter Stress-Induced Patterns of Global DNA Methylation in Brain and Blood. Molecular Nutrition & Food Research. 2018;62:1700722.

44. Kelsey JE, Carlezon WA, Falls WA. Lesions of the nucleus accumbens in rats reduce opiate reward but do not alter context-specific opiate tolerance. Behav Neurosci. 1989;103:1327–1334.

45. Frolinger T, Sims S, Smith C, Wang J, Cheng H, Faith J, et al. The gut microbiota composition affects dietary polyphenols-mediated cognitive resilience in mice by modulating the bioavailability of phenolic acids. Sci Rep. 2019;9:3546.

46. Westfall S, Pasinetti GM. The Gut Microbiota Links Dietary Polyphenols With Management of Psychiatric Mood Disorders. Frontiers in Neuroscience. 2019;13.

47. Hills RD, Pontefract BA, Mishcon HR, Black CA, Sutton SC, Theberge CR. Gut Microbiome: Profound Implications for Diet and Disease. Nutrients. 2019;11.

48. Fields CT, Sampson TR, Bruce-Keller AJ, Kiraly DD, Hsiao EY, de Vries GJ. Defining Dysbiosis in Disorders of Movement and Motivation. J Neurosci. 2018;38:9414–9422.

49. Anhê Ff, Roy D, Pilon G, Dudonné S, Matamoros S, Varin TV, et al. A polyphenol-rich cranberry extract protects from diet-induced obesity, insulin resistance and intestinal inflammation in association with increased Akkermansia spp. population in the gut microbiota of mice. Gut. 2015;64:872–883.

50. Roopchand DE, Carmody RN, Kuhn P, Moskal K, Rojas-Silva P, Turnbaugh PJ, et al. Dietary Polyphenols Promote Growth of the Gut Bacterium Akkermansia muciniphila and Attenuate High-Fat Diet-Induced Metabolic Syndrome. Diabetes. 2015;64:2847–2858.

51. Segata N, Waldron L, Ballarini A, Narasimhan V, Jousson O, Huttenhower C. Metagenomic microbial community profiling using unique clade-specific marker genes. Nat Methods. 2012;9:811–814.

52. Ferguson D, Koo JW, Feng J, Heller E, Rabkin J, Heshmati M, et al. Essential role of SIRT1 signaling in the nucleus accumbens in cocaine and morphine action. J Neurosci. 2013;33:16088–16098.

53. Han Y, Jiang C, Tang J, Wang C, Wu P, Zhang G, et al. Resveratrol reduces morphine tolerance by inhibiting microglial activation via AMPK signalling. Eur J Pain. 2014;18:1458–1470.

54. He X, Ou P, Wu K, Huang C, Wang Y, Yu Z, et al. Resveratrol attenuates morphine antinociceptive tolerance via SIRT1 regulation in the rat spinal cord. Neurosci Lett. 2014;566:55–60.

55. Tsai R-Y, Wang J-C, Chou K-Y, Wong C-S, Cherng C-H. Resveratrol reverses morphine-induced neuroinflammation in morphine-tolerant rats by reversal HDAC1 expression. J Formos Med Assoc. 2016;115:445–454.

56. Tsai R-Y, Chou K-Y, Shen C-H, Chien C-C, Tsai W-Y, Huang Y-N, et al. Resveratrol regulates N-methyl-D-aspartate receptor expression and suppresses neuroinflammation in morphine-tolerant rats. Anesth Analg. 2012;115:944–952.

57. Renthal W, Kumar A, Xiao G, Wilkinson M, Covington HE, Maze I, et al. Genome-wide analysis of chromatin regulation by cocaine reveals a role for sirtuins. Neuron. 2009;62:335–348.

58. Li Y, Yu L, Zhao L, Zeng F, Liu Q. Resveratrol modulates cocaine-induced inhibitory synaptic plasticity in VTA dopamine neurons by inhibiting phosphodiesterases (PDEs). Sci Rep. 2017;7:15657.

59. Morris G, Gamage E, Travica N, Berk M, Jacka FN, O’Neil A, et al. Polyphenols as adjunctive treatments in psychiatric and neurodegenerative disorders: Efficacy, mechanisms of action, and factors influencing inter-individual response. Free Radical Biology and Medicine. 2021;172:101–122.

60. Paluszczak J, Krajka-Kuźniak V, Baer-Dubowska W. The effect of dietary polyphenols on the epigenetic regulation of gene expression in MCF7 breast cancer cells. Toxicol Lett. 2010;192:119–125.

61. Chung S, Yao H, Caito S, Hwang J-W, Arunachalam G, Rahman I. Regulation of SIRT1 in cellular functions: role of polyphenols. Arch Biochem Biophys. 2010;501:79–90.

## Supplementary References

1. Torre D, Lachmann A, Ma’ayan A. BioJupies: Automated Generation of Interactive Notebooks for RNA-Seq Data Analysis in the Cloud. Cell Syst. 2018;7:556-561.e3.

2. Xia J, Gill EE, Hancock REW. NetworkAnalyst for statistical, visual and network-based meta-analysis of gene expression data. Nat Protoc. 2015;10:823–844.

3. Zhou G, Soufan O, Ewald J, Hancock REW, Basu N, Xia J. NetworkAnalyst 3.0: a visual analytics platform for comprehensive gene expression profiling and meta-analysis. Nucleic Acids Res. 2019;47:W234–W241.

4. Raudvere U, Kolberg L, Kuzmin I, Arak T, Adler P, Peterson H, et al. g:Profiler: a web server for functional enrichment analysis and conversions of gene lists (2019 update). Nucleic Acids Res. 2019;47:W191–W198.

5. Chen EY, Tan CM, Kou Y, Duan Q, Wang Z, Meirelles GV, et al. Enrichr: interactive and collaborative HTML5 gene list enrichment analysis tool. BMC Bioinformatics. 2013;14:128.

6. Kuleshov MV, Jones MR, Rouillard AD, Fernandez NF, Duan Q, Wang Z, et al. Enrichr: a comprehensive gene set enrichment analysis web server 2016 update. Nucleic Acids Res. 2016;44:W90–W97.

7. Magoč T, Salzberg SL. FLASH: fast length adjustment of short reads to improve genome assemblies. Bioinformatics. 2011;27:2957–2963.

8. Caporaso JG, Kuczynski J, Stombaugh J, Bittinger K, Bushman FD, Costello EK, et al. QIIME allows analysis of high-throughput community sequencing data. Nat Methods. 2010;7:335–336.

9. Douglas GM, Maffei VJ, Zaneveld JR, Yurgel SN, Brown JR, Taylor CM, et al. PICRUSt2 for prediction of metagenome functions. Nat Biotechnol. 2020;38:685–688.

10. Simpson GR, Riley AL. Morphine preexposure facilitates morphine place preference and attenuates morphine taste aversion. Pharmacol Biochem Behav. 2005;80:471–479.

11. Zhao W, Wang J, Bi W, Ferruzzi M, Yemul S, Freire D, et al. Novel application of brain-targeting polyphenol compounds in sleep deprivation-induced cognitive dysfunction. Neurochem Int. 2015;89:191–197.

12. Ho L, Ferruzzi MG, Janle EM, Wang J, Gong B, Chen T-Y, et al. Identification of brain-targeted bioactive dietary quercetin-3-O-glucuronide as a novel intervention for Alzheimer’s disease. FASEB J Off Publ Fed Am Soc Exp Biol. 2013;27:769–781.

13. Wang J, Ferruzzi MG, Ho L, Blount J, Janle EM, Gong B, et al. Brain-targeted proanthocyanidin metabolites for Alzheimer’s disease treatment. J Neurosci Off J Soc Neurosci. 2012;32:5144–5150.

14. Wang J, Bi W, Cheng A, Freire D, Vempati P, Zhao W, et al. Targeting multiple pathogenic mechanisms with polyphenols for the treatment of Alzheimer’s disease-experimental approach and therapeutic implications. Front Aging Neurosci. 2014;6:42.

15. Jiang C, Sakakibara E, Lin W-J, Wang J, Pasinetti GM, Salton SR. Grape-derived polyphenols produce antidepressant effects via VGF- and BDNF-dependent mechanisms. Ann N Y Acad Sci. 2019;1455:196–205.

16. Jiang C, Lin W-J, Sadahiro M, Labonté B, Menard C, Pfau ML, et al. VGF function in depression and antidepressant efficacy. Mol Psychiatry. 2018;23:1632–1642.

17. Wang J, Hodes GE, Zhang H, Zhang S, Zhao W, Golden SA, et al. Epigenetic modulation of inflammation and synaptic plasticity promotes resilience against stress in mice. Nat Commun. 2018;9:477.

18. Frolinger T, Sims S, Smith C, Wang J, Cheng H, Faith J, et al. The gut microbiota composition affects dietary polyphenols-mediated cognitive resilience in mice by modulating the bioavailability of phenolic acids. Sci Rep. 2019;9:3546.

19. Dalile B, Van Oudenhove L, Vervliet B, Verbeke K. The role of short-chain fatty acids in microbiota-gut-brain communication. Nat Rev Gastroenterol Hepatol. 2019;16:461–478.

20. Hofford RS, Mervosh NL, Euston TJ, Meckel KR, Orr AT, Kiraly DD. Alterations in microbiome composition and metabolic byproducts drive behavioral and transcriptional responses to morphine. Neuropsychopharmacol Off Publ Am Coll Neuropsychopharmacol. 2021. 14 June 2021. https://doi.org/10.1038/s41386-021-01043-0.

21. Kiraly DD, Walker DM, Calipari ES, Labonte B, Issler O, Pena CJ, et al. Alterations of the Host Microbiome Affect Behavioral Responses to Cocaine. Sci Rep. 2016;6:35455.

22. Barbieri R, Coppo E, Marchese A, Daglia M, Sobarzo-Sánchez E, Nabavi SF, et al. Phytochemicals for human disease: An update on plant-derived compounds antibacterial activity. Microbiol Res. 2017;196:44–68.

